# PSMD14 drives melanoma cell survival and MAPK inhibitor resistance through histone H2A deubiquitination

**DOI:** 10.64898/2026.02.27.708453

**Authors:** Michael Ohanna, Pierric Biber, Mira Kahil, Serena Diazzi, Lauren Lefevre, Frédéric Larbret, Robin Didier, Stéphane Audebert, Véronique Delmas, Lionel Larue, Sophie Tartare-Deckert, Marcel Deckert

## Abstract

Melanoma progression and resistance to targeted therapies remain major clinical challenges, driven in part by the remarkable phenotypic plasticity of melanoma cells. Identifying molecular mechanisms that couple tumor survival to adaptative drug responses is therefore essential. Here, we identify PSMD14, a proteasome-associated deubiquitinase, as an essential regulator of melanoma plasticity, growth, survival, and therapeutic resistance with strong prognostic significance in metastatic disease. An unbiased siRNA screen targeting the human deubiquitinase family revealed PSMD14 (proteasome 26S subunit, non-ATPase 14) as a top regulator of melanoma cell proliferation. Integrative analyses of DepMap, TCGA, and patient-derived datasets revealed that PSMD14 is frequently upregulated in melanoma, enriched in metastatic lesions, and significantly associated with poor patient outcome. Functional and pharmacological studies demonstrated that genetic depletion or inhibition of PSMD14 suppresses proliferation, clonogenic and long-term growth, and viability of melanoma cells across BRAF-, NRAS-, and NF1-driven genotypes, while inducing DNA damage and apoptosis. Consistently, PSMD14 inhibition markedly reduced tumor growth in Nras and Braf syngeneic mouse models. Mechanistically, we uncover a non-proteolytic role for PSMD14 as an epigenetic regulator of chromatin state. Proteomic and biochemical analyses identified histone H2A as a direct interactor and substrate of PSMD14. PSMD14 deubiquitinates H2A at lysine 119 independently of the proteasome, antagonizing the Polycomb E3 ligase RING1B. Loss of PSMD14 allows the increment of H2AK119 ubiquitination, transcriptional repression of pro-survival genes, including MCL1 and BCL2, and apoptotic cell death, effects rescued by RING1B depletion. Importantly, we demonstrate that the PSMD14-H2A axis governs melanoma adaptation to MAPK pathway inhibition. PSMD14 expression and H2AK119 ubiquitination dynamically correlate with therapeutic response, drug-tolerant persistence, and acquired resistance. Targeting PSMD14 genetically or pharmacologically enhances the efficacy of BRAF and MEK inhibitors, suppresses the emergence of drug-tolerant persister cells, and prevents tumor relapse *in vivo*. Together, this findings establish PSMD14 as a chromatin-rewiring enzyme that links proteostasis to epigenetic control of melanoma plasticity and therapy resistance, highlighting PSMD14 as a promising biomarker and therapeutic target in aggressive and drug-resistant melanoma.

## INTRODUCTION

A defining hallmark of malignant cells is their remarkable phenotypic plasticity, allowing them to adapt dynamically to microenvironmental cues and therapeutic stress. This adaptative capacity drives treatment evasion and the emergence of drug resistance ^1^. While genetic alterations and transcriptional reprogramming are well established drivers of cancer plasticity, tumor cells also depend on dynamic and extensive remodeling of protein homeostasis to sustain oncogenic growth and therapy resistance ^2^. However, compared with genetic and epigenetic mechanisms, the contribution of proteostasis adaptation to cancer cell plasticity remains poorly understood.

Cutaneous melanoma represents a paradigmatic example of a highly plastic and therapy-resistant malignancy. Most melanomas are driven by oncogenic mutations in BRAF and NRAS ^3,4^, which constitutively engage the MAPK signaling pathway and stimulate transcriptional programs that promotes cell proliferation and survival ^3,5^. These discoveries let to the development of targeted therapies, notably BRAF^V600E^ and MEK inhibitors (BRAFi/MEKi), which have markedly improved patient outcomes. Nevertheless, therapeutic response are invariably transient, as innate or acquired resistance and disease relapse almost universally occur ^6^. Mounting evidence indicates that resistance to MAPK pathway inhibition is largely attributable to intrinsic melanoma cell plasticity and phenotype switching ^7–9^, involving a complex interplay between genetic and epigenetic alterations ^10–13^ and adaptative rewiring of signaling pathways ^10,14–16^. Moreover, recent findings suggest that melanoma cells must actively remodel pathways controlling the proteome to maintain survival, plasticity, and adaptability under oncogenic and therapeutic stress ^17–20^.

Protein homeostasis is centrally regulated by the ubiquitin–proteasome system (UPS) ^21^. Ubiquitination is a versatile and key post-translational modification that regulates protein stability, localization, and activity through the covalent attachment of ubiquitin to lysine residues on target proteins via a hierarchical E1-E2-E3 ligase cascade ^22^. The functional outcome depends on the nature of ubiquitin modification : monoubiquitination typically regulates non-proteolytic processes such as DNA repair, endocytosis, and transcriptional activation, whereas polyubiquitination, particularly via K48- or K11-linked chains, typically targets proteins for proteasomal degradation ^22^. This ubiquitin signals are dynamically reversed or edited by deubiquitinases (DUBs), a family of more than 90 enzymes that precisely regulate ubiquitin-dependent processes chains ^23^. Aberrant expression, mutation, or dysregulation of DUBs has been implicated in tumorigenesis, highlighting their potential as therapeutic targets ^24,25^. Several DUBs have been directly linked to melanoma progression and therapeutic response. For instance, USP9X promotes tumorogenicity through ETS1 in NRAS-mutant melanoma ^26^ and contribute to mechanical adaptation to targeted therapies in BRAF-mutant melanoma ^20^. Targeting USP14, a proteasome-associated DUB, triggers caspase-independent cell death and overcomes resistance to to MAPK inhibitors ^27^. Conversely, down-regulation of CYLD promotes melanoma progression ^28^, whereas reduced USP7 expression favors senescence in both BRAF- and NRAS-mutant melanoma ^29^. Despite these insights, the broader contribution of DUBs to melanoma plasticity and therapeutic adaptation remains largely unexplored.

Here, we identified PSMD14 (also known as RPN11 or POH1), a proteasome-associated deubiquitinase, as a central regulator of melanoma survival, tumor growth, and resistance to targeted therapies. Using an unbiased siRNA screen, we uncover PSMD14 as an essential determinant of melanoma cell viability across distinct oncogenic backgrounds. Genetic depletion or pharmacological inhibition of PSMD14 suppresses proliferation, clonogenic growth, and viability, induces DNA damage, and impairs tumor growth in both BRAF- and NRAS-driven melanoma models. Mechanistically, we revealed an unexpected, non-proteolytic function of PSMD4 as an epigeneric regulator of chromatin state. Proteomic and biochemical analyses identified histone variant H2A as a previously unrecognized direct interactor and functional substrate of PSMD14 in melanoma. We demonstrate that PSMD14 deubiquitinated histone H2A at lysine 119 independently of the proteasome, antagonizing the Polycomb E3 ubiquitin ligase RING1B. Loss or inhibition of PSMD14 increased H2AK119 ubiquitination (H2AK119ub), transcriptional repression of prosurvival genes, including the anti-apoptotic factors *BCL2* and *MCL-1*, and induction of melanoma cell death. These effects were rescued by RING1B depletion, establishing a functional antagonism between PSMD14 and Polycomb-mediated chromatin repression. Given the central role of histone post-translational modifications as non-genetic determinants of therapeutic response, we further investigated the role of PSMD14-H2A axis in melanoma adaptation to MAPK pathway inhibition. We showed that MAPK inhibition dynamically modulates PSMD14 expression and H2AK119 ubiquitination, and that targeting this axis synergized with BRAF and MEK inhibitors to prevent the emergence of drug-tolerant and drug-resistant melanoma cells both *in vitro* and *in vivo*. Altogether, our findings uncover PSMD14 as a chromatin-rewiring enzyme that links proteostasis to epigenetic regulator of melanoma plasticity and drug therapy resistance, highlighting PSMD14 as a promising therapeutic target in metastatic and drug-resistant melanoma.

## RESULTS

### Identification of *PSMD14* as an essential gene linked to poor prognosis in metastatic melanoma

We performed an unbiased genetic screen to identify deubiquitinating enzymes (DUBs) involved in melanoma cell proliferation and survival. The screen was conducted in 501Mel cells a well-characterized BRAF mutant melanoma cell line harboring a proliferative phenotype ^7^. 501Mel cells were transfected with a human DUB siRNA library consisting of pools of four oligonucleotide sequences targeting 98 human DUBs (Dharmacon siGENOME® SMARTpool® siRNA Library-Human Deubiquitinating Enzymes), along with non-targeting siRNAs controls. Cell proliferation was followed for 96 h by live-cell imaging-based measurement of cell confluence **(Figure 1A)**. The effects of individual siDUBs pools were then ranked according to their impact on cell confluency at 96 h compared to the mean of non-targeting control siRNA conditions **(Figure 1B, Supplementary Table 1)**. Importantly, our screen confirmed the importance of USP14 in regulating melanoma cell proliferation **(Supplementary Table 1),** consistent with our previous studies (Didier et al., 2018), thereby validating the robustness of our approach. In addition, knockdown of seven other DUBs resulted in a severe inhibition of 501Mel cell proliferation after 96 h with more than 50% inhibition of cell confluency. To further prioritize candidates, we interrogated the Achilles dependency data set (DepMap Public 20Q3), which provides a genome-wide CRISPR-Cas9 knockout screens. Among the seven DUBs identified in our screen, PSMD14 (also known as RPN11 or POH1) exhibited the most negative CERES score in skin cancer, further supporting its role as an essential gene for cell viability in melanoma cells **(Figure 1C)**. In addition to melanoma, PSMD14 dependency was also observed across a broad range of solid and blood cancers **(Figure 1D)**. PSMD14 is a Zn2+-dependent metalloisopeptidase of the JAMM family and an intrinsic component of the 19S regulatory particle of the proteasome that plays a critical role in proteasomal function and tumour growth ^30–33^. Based on these observations, we hypothesized that PSMD14 constitutes a key contributor of melanoma biology and therapeutic response. Analysis of GEPIA databases revealed that PSMD14 mRNA expression is significantly upregulated in melanoma tissue samples compared with normal tissue samples **(Figure 1E)**. Moreover, PSMD14 expression correlated with melanoma progression, with higher levels detected in metastatic melanoma relative to benign skin lesions (nevi) **(Figure 1F)**. Consistent with these findings, TCGA melanoma cohort analysis revealed that PSMD14 is altered in 12% of melanoma of cases, predominantly through increased RNA expression **(Supplementary Figure 1A)**. High PSMD14 levels was significantly associated with poor outcomes in melanoma patients (P=0.008) **(Figure 1G)**. Gene set enrichment analysis (GSEA) of TCGA melanoma transcriptomic data revealed a positive correlation between PSMD14 levels and key gene signature pathways involved in tumor growth and drug resistance, including cell cycle and proliferation, PI3K/AKT/mTOR, and metabolic pathways **(Figure 1H, Supplementary Figure 1B, C)**. Collectively, these findings suggest that PSMD14 plays a crucial role in melanoma cell viability, disease progression, and patient outcome, highlighting its potential as both a biomarker for melanoma staging and a therapeutic target.

**Figure 1.**
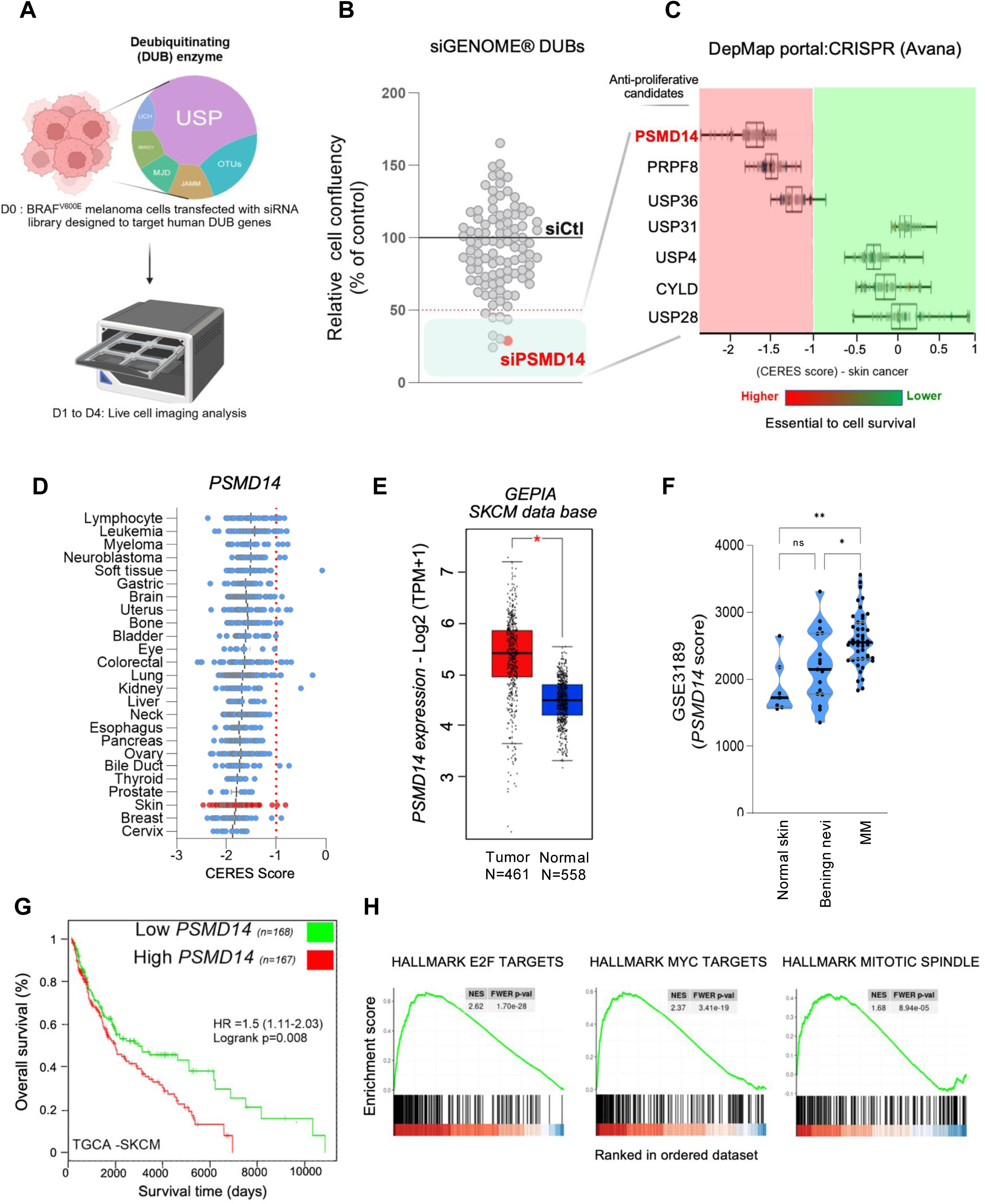
PSMD14 is an essential gene linked to poor prognosis in metastatic melanoma. **(A)** Schematic diagram illustrating the strategy to identify deubiquitinases (DUBs) affecting proliferation and survival in BRAF mutant melanoma cells. siRNA knockdown in 501Mel cells was used, with confluence measured by IncuCyte at 96 h and results normalized to a non-targeting control. **(B)** Dot plots show normalized cell confluency for each siRNA versus siCtl. Candidates within the blue square demonstrate over 50% inhibition of confluence. PSMD14, highlighted in red, appears as one of the top hits. **(C)** Bar graph showing the correlation between dependency scores and candidates from siRNA DUB screening in skin cancer cell lines, using DepMap CRISPR data. Significant dependencies < -1 (red panel) and > -1 (green panel) illustrate essential genes and non-essential genes, respectivey. *PSMD14* shows the strongest impact on cell survival. **(D)** Dependency scores of *PSMD14* across pan-cancer cell lines, based on DepMap CRISPR data (21Q4). Melanoma cell lines, highlighted in red, are among the most affected by PSMD14 depletion. **(E)** *PSMD14* expression levels in primary melanoma tumors and normal skin tissues, using GEPIA interactive analysis. *, *P*<0.05. (**F)** PSMD14 expression across melanoma progression stages (GSE3189). **(G)** Kaplan-Meier overall survival curves in melanoma patients with high or low PSMD14 expression from TGCA SKCM dataset were obtained through SurvExpress (p=0.008; log-rank test). **(H)** GSEA of the TCGA SKCM dataset shows enrichment of hallmark gene sets including E2F targets, MYC targets and mitotic spindle in PSMD14^high tumors (positive NES; significant FWER-adjusted p-values)

### Targeting PSMD14 impairs melanoma cell proliferation and survival *in vitro* and *in vivo*

To further investigate the role of PSMD14 in melanoma, we knocked down PSMD14 in BRAF mutant cell lines 501Mel and A375 using three independent siRNA sequences and monitored cell proliferation by live-cell imaging. Consistent with the results of our screen, all three siPSMD14 sequences severely reduced melanoma cell proliferation after 30h to 48h of transfection **(Figure 2A)**. PSMD14 depletion was associated with the downregulation of the cell-cycle dependent kinase CDK2 and a strong induction of the CDK inhibitor p21Cip1 **(Figure 2B)**, indicating cell cycle arrest. Since clonogenic growth is a hallmark of malignant cells, we evaluated the impact of PSDM14 on colony formation. Compared with control siRNA, UACC62, A375 and 501Mel cells transfected with PSMD14 siRNAs were unable to form colonies after 7 days, showing that PSMD14 expression is required for melanoma clonogenic growth **(Supplementary Figure 2A, B).** We next examined the role of PSMD14 in cell survival across a collection of melanoma cell lines with diverse mutational backgrounds, transcriptional states, and metastatic potential, representative of three major melanoma subtypes defined by BRAF, NRAS and NF1 mutations ^34^. PSMD14 was knocked down in seven melanoma cell lines harboring activating mutations in BRAF (WM793, WM9, G361), NRAS (Sbcl2, WM2032), or hemizygous deletion of NF1 (MEWO), as well as in two short term cultures derived from BRAF^V600E^ metastatic melanoma (MM029 and MM099) **(Figure 2C)**. In all models examined, PSMD14 depletion strongly reduced melanoma cell viability regardless of the type of oncogenic driver mutation, and regardless of their transcriptional cell state, or metastatic origine. Conversely, overexpression of tagged PSMD14 in A375 cells **(Figure 2D)** significantly increases long-term cell proliferation **(Figure 2E, Supplementary Figure 2C, D)**. In contrast, overexpression of a catalytic inactive PSMD14 mutant harboring alanine substitutions of two conserved histidines in the MPN/JAMM catalytic site ^31^ (PSMD14 His113/115Ala, hereafter named PSMD14 JAMM^M^) **(Figure 2D)** impaired melanoma cell growth **(Figure 2E, Supplementary Figure 2C, D)**. These results show that the deubiquitinating activity of PSMD14 is required to sustain melanoma cell proliferation. To pharmacologically validate these findings, we treated melanoma cells with 8-TQ, a recently described PSMD14 inhibitor (PSMD14i) that binds the JAMM catalytic domain ^32^. PSMD14i induced dose-dependent growth with comparable IC50 values (0.25 to 0.75μM) across multiple human melanoma cell lines (MM099, MM029, Sbcl2, 501Mel, WM9 and MEWO and mouse melanoma lines (YUMM1.7 and MaNRAS) **(Figure 2F)**. We further observed that PSMD14 inhibition or depletion induces apoptosis, as evidenced by increased Annexin V/PI cell staining **(Figure 2G, H)**, along with caspase 3 and PARP cleavage **(Figure 2I)**. At a molecular level, PSMD14 inhibition or depletion resulted in a massive accumulation of polyubiquitylated proteins **(Supplementary Figure 2E, F)**, and activation of a DNA damage response, indicated by increased phosphorylation of nuclear histone H2AX (pH2AX) **(Figure 2J and K, Supplementary Figure 2G)**.

**Figure 2.**
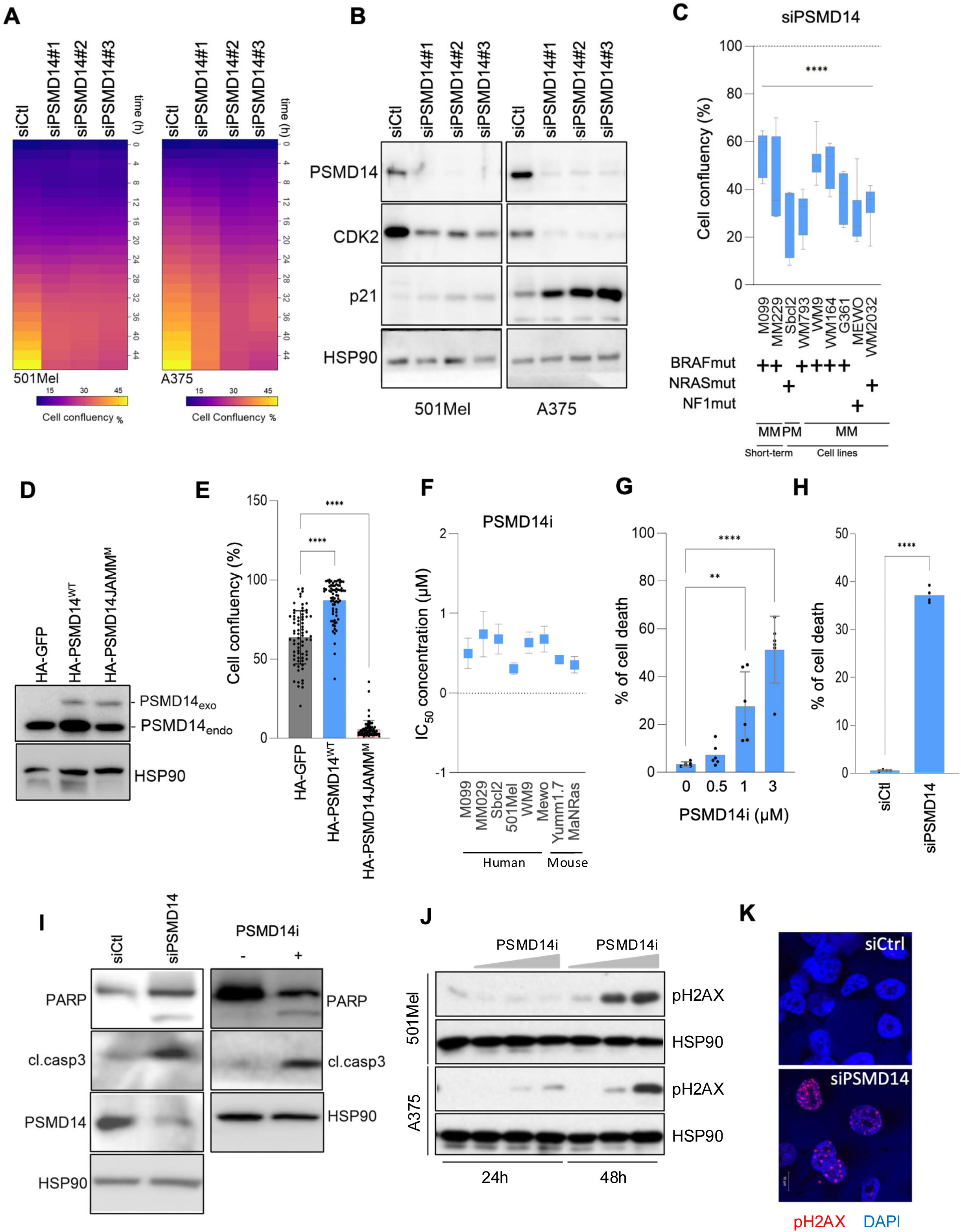
Inhibition of PSMD14 suppresses BRAF- and NRAS-mutant melanoma proliferation by inducing apoptosis. **(A)** Time-course heatmap showing cell confluency following PSMD14 knockdown with three independent siRNA sequences (siPSMD14#1, #2, and #3) compared to control siRNA (siCtl) in 501Mel and A375 BRAF-mutant melanoma cells. **(B)** Western blot analysis of PSMD14, CDK2, and p21Cip1 expression in 501Mel and A375 melanoma cells transfected with control siRNA (siCtl) or siPSMD14#1, #2, and #3. HSP90, loading control. **(C)** Quantification of cell survival after 72 h of PSMD14 depletion in melanoma cell lines harboring distinct oncogenic mutations. Data are mean ± SEM (n=6). \*\*\*\**P*<0.0001, Wilcoxon signed-rank test. **(D)** Western blot analysis with anti-PSMD14 antibody showing expression of exogenous HA-PSMD14 and HA-PSMD14 JAMM^M^ compared to endogenous PSMD14 in A375 cells. HSP90, loading control. **(E)** Bar graph showing cell confluency of HA-PSMD14, HA-PSMD14 JAMM^M^ and HA-GFP-expressing A375 cells after 72 h. Data are presented as mean ± SEM (n = 3). \*\*\*\**P*<0.0001, one-way ANOVA. **(F)** IC_50_ determination after dose response of PSMD14i (8TQ) on human and murine melanoma cell lines with various oncogenic mutations. The corresponding IC_50_ were determined by measuring the cell viability following AnnexinV and DAPI staining and by flow cytometry analysis at 72 h (mean ± SEM, n=3, two-way ANOVA). **(G)** Flow cytometry analysis of apoptosis after 72 h of PSMD14i treatment in A375 cells using Annexin V-FITC and DAPI staining. Bar graphs display the percentage of Annexin V- and DAPI-positive cells. Data are mean ± SEM (n = 6), \*\**P*<0.01, \*\*\*\**P*<0.0001, two-way ANOVA. **(H)** Flow cytometry analysis of apoptosis on A375 cells transfected for 72h with siCtl or siPSMD14. Bar graphs display the percentage of Annexin V- and DAPI-positive cells (mean ± SEM, n = 3, \*\*\*\**P*<0.0001, two-way ANOVA. **(I)** Western blot analysis of apoptosis markers following PSMD14 inhibition or knockdown. HSP90, loading control. **(J)** Western blot analysis of gH2AX expression in a dose- and time-dependent manner following PSMD14i treatment in A375 and 501Mel cells. **(K)** Confocal microscopy analysis shows a marked increase in γH2AX-rich nuclear foci in PSMD14-depleted (siPSMD14, 75 ± 10 foci per nucleus) compared to control (siCtl, 5 ± 3 foci per nucleus) A375 cells with γH2AX foci in red and DAPI-stained nuclei in blue. Scale bar, 10 µm.

Next, we assessed the anti-tumor activity of PSMD14i in syngeneic mouse models of of BRAF- and NRAS-mutant melanoma. Established tumors derived from YUMM1.7 and MaNRAS cells were treated with PSMD14i, and tumor development was monitored over time. PSMD14 inhibition significant suppressed tumor growth in both genetic backgrounds **(Figure 3A)**. Ex vivo analysis of YUMM1.7 tumors further showed that PSMD14 targeting reduced tumor proliferation and increased DNA damage and apoptosis, as shown by decreased Ki-67 staining and cleaved caspase 3 levels, respectively **(Figure 3B)**. Collectively, these results establish PSMD14 as a critical regulator of melanoma cell proliferation and survival *in vitro* and *in vivo*, highlighting its potential as a therapeutic vulnerability across melanoma subtypes.

**Figure 3.**
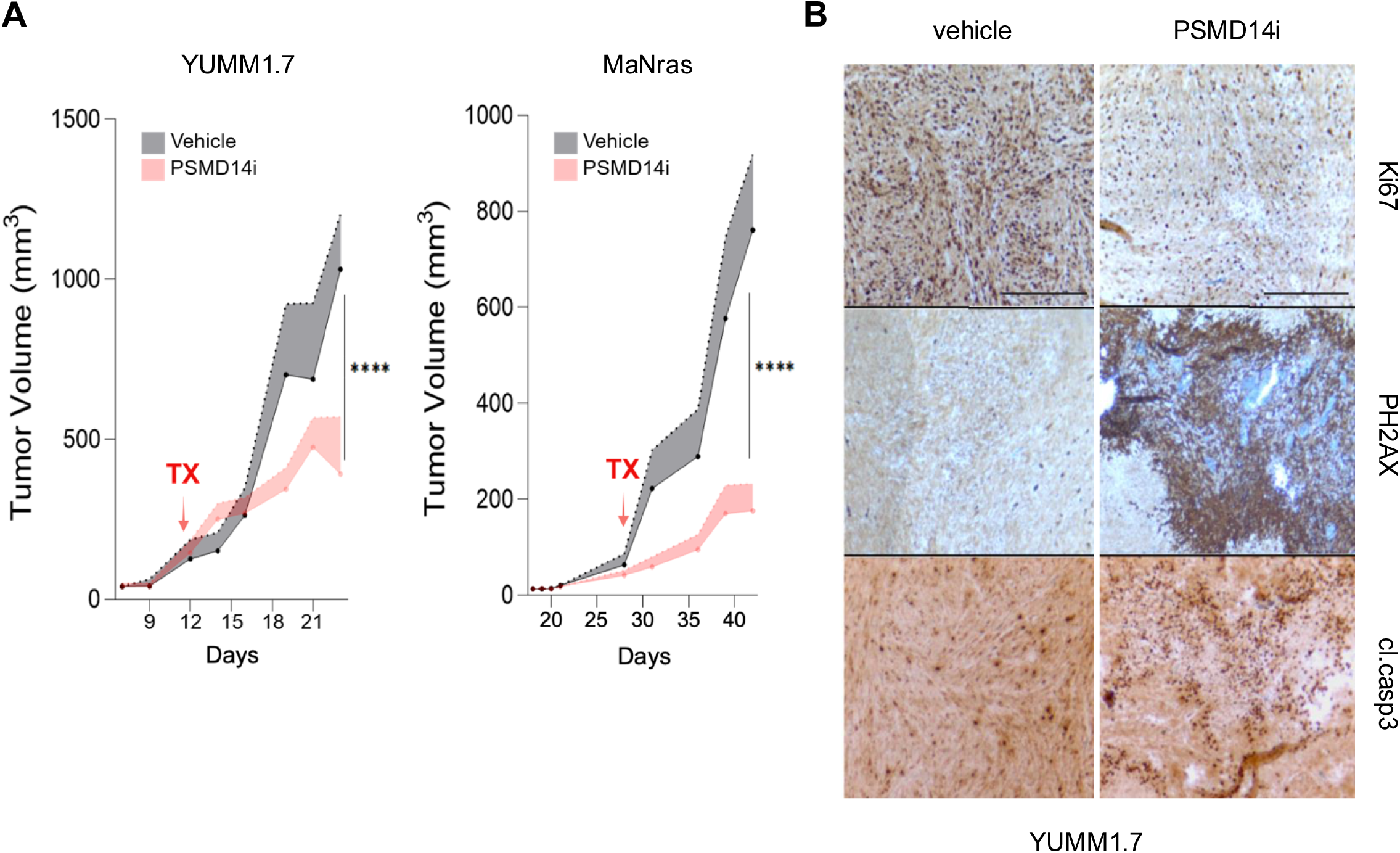
Inhibition of PSMD14 reduces tumor growth in BRAFV600E- and NRAS-mutant melanoma models. **(A, B)** Effect of PSMD14 inhibitor (8TQ; 15 mg/kg) on tumor growth in **(A)** YUMM1.7 murine melanoma (Braf^V600E^/Pten^null^/Cdkn2a^null^) and **(B)** MaNRAS melanoma (Nras^Q61K^) models. Tumor volumes were measured over time in mice treated with vehicle (n = 10) or PSMD14i (n = 8). Statistical significance was determined using two-way repeated-measures ANOVA followed by Bonferroni’s multiple comparisons test (**P < 0.01; ***P < 0.001). **(C)** Representative immunohistochemical staining of Ki67, γH2AX, and cleaved caspase-3 in YUMM1.7 tumors following 2 weeks of treatment with vehicle or PSMD14i. Scale bar, 100 µm.

### PSMD14 directly deubiquitinates histone H2A to regulate chromatin-associated processes

To unravel the mechanisms governing PSMD14’s effects on melanoma, we sought to identify its interaction partners and explore potential non proteasomal roles. We performed immunoaffinity purification of HA-tagged PSMD14 from A375 cells stably expressing HA-PSMD14, followed by mass spectrometry-based proteomic analysis of the immunoprecipitates **(Figure 4A)**. Across three independent experiments, we identified 705 high-confidence PSMD14 interacting proteins, each represented by at least one unique peptide and benchmarked against HA-GFP **(Figure 4B)**. Gene Ontology (GO) enrichment analysis revealed a strong over-representation of biological processes related to protein deubiquitination related to protein deubiquitination (GO:0016579) and proteasome assembly (GO:0043248) **(Figure 4C)**, consistent with the described function of PSMD14 as a proteasome-associated deubiquitinase ^30,31^. Notably, additional enriched categories were associated with chromatin remodeling and DNA damage repair, including chromatin silencing (GO:0006342), double strand break repair via non-homologous end joining (GO:0006303), and epigenetic regulation of gene expression (GO:0040029). Among the significantly enriched interactors were multiple histone proteins, including variants of histone H2A **(Figure 4B)**. Histone ubiquitination is a key epigenetic mechanism regulating transcription and DNA damage repair, and mono-ubiquitination of histone H2A at lysine 119 (H2AK119ub) has been implicated in tumorigenesis and melanoma progression ^35,36^. This led us to hypothesize that PSMD14 directly interacts with the histone H2A variants and promotes its deubiquitination potentially counteracting the activity of the Polycomb E3 ubiquitin ligase RING1B/RNF2, which catalyzes H2AK119 monoubiquitination and mediates transcriptional repression **(Figure 4D)**. To validate the interaction between PSMD14 and H2A, we performed co-immunoprecipitation assays in A375 HA-PSMD14 cells and confirmed the association of PSMD14 with H2A, alongside known PSMD14-binding proteins, such as, PSMD12 and PSMD7 **(Figure 4E)**. Further, HA-based pulldown assays using Flag-tagged H2A and HA-tagged PSMD14 in HEK cells confirmed the direct interaction between PSMD14 and H2A **(Figure 4F)**.

**Figure 4.**
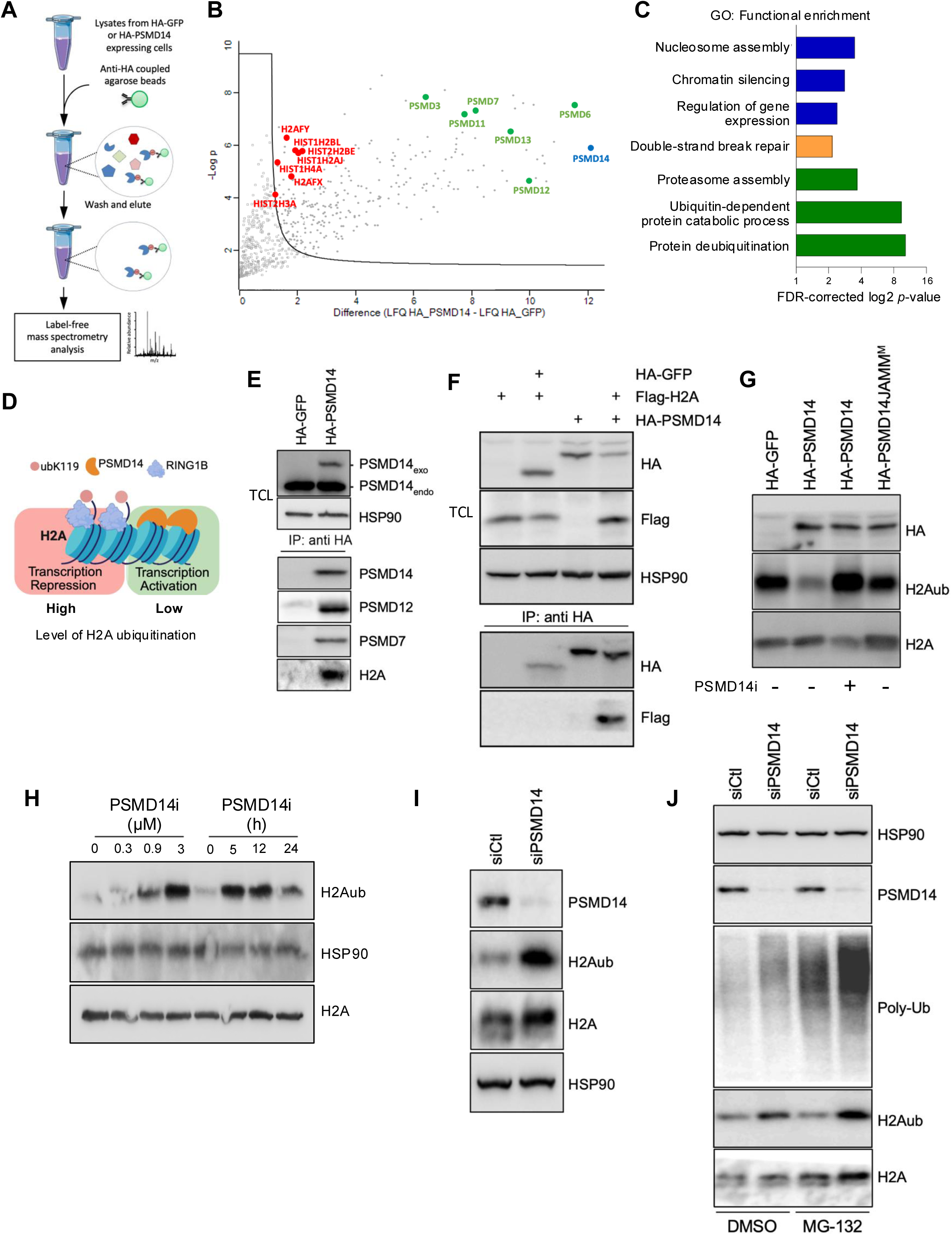
PSMD14 interacts with and deubiquitinates histone H2A. **(A)** Schematic illustration of the strategy to study the PSMD14 interactome in melanoma cells. Lysates from A375 expressing HA-GFP or HA-PSMD14 cells are immunoprecipitated using anti-HA coated agarose beads and analyzed by quantitative mass spectrometry. **(B)** Volcano plot showing significantly differential enrichment of PSMD14 interactors over GFP. Relative intensity-based label-free quantification (LFQ) was processed using the MaxLFQ algorithm. A two-sample *t*-test was performed using permutation-based FDR-controlled at 0.5 %. The *p-*value was adjusted using a scaling factor S0 with a value of 1. The difference LFQ intensity was plotted against the −log10 of the *p-*value. The bold line indicated the applied threshold values (*p-*value < 0.05; fold change ≥ 1.5). Among PSMD14 (blue dot), components of the proteasome (green dots) and histones (red dots) are highlighted. **(C)** Bar graph showing gene ontology (GO) analysis of the identified PSMD14-interacting proteins grouped into functional categories. **(D)** Schematic representation of the proposed function of PSMD14 in regulating H2A ubiquitination (H2Aub) and relieving H2Aub-dependent transcriptional repression. **(E)** Co-immunoprecipitation analysis of HA-PSMD14 confirming the binding of PSMD14 to proteasome-associated proteins PSMD7 and PSMD12, to histone H2A in A375 cells. **(F)** Co-immunoprecipitation of HA–PSMD14 and FLAG–H2A overexpressed in HEK293T cells. **(G)** Western blot analysis of H2Aub and total H2A levels following overexpression of HA–PSMD14 alone or in combination with PSMD14 inhibitor (PSMD14i) or following overexpression of HA-PSMD14 JAMM^M^ mutant in HEK293T cells. **(H)** Western blot analysis of H2Aub levels in A375 melanoma cells treated with increasing concentrations of PSMD14i for 24h or over time with 1µM of PSMD14i. **(I)** Western blot detection of elevated H2Aub levels in A375 cells after 72 h of PSMD14 depletion (siPSMD14) compared to control (siCtl). HSP90, loading control. **(J)** Western blot analysis of A375 cells transfected with or without PSMD14 siRNA and subsequently treated with MG132 (20 μg/mL for 4 h), using the indicated antibodies. HSP90, loading control.

We next investigated whether PSMD14 regulates H2A ubiquitination through its catalytic activity. In HEK cells, expression of wild-type PSMD14 resulted in a marked reduction of H2Aub levels, whereas expression of catalytically inactive JAMM^M^ mutant failed to do so **(Figure 4G)**. Consistent with these findings, melanoma cells exposed to PSMD14i **(Figure 4H)** or knocked down of PSMD14 by siRNA **(Figure 4I)** displayed a significant accumulation of H2A-K119ub, while total H2A levels remained unchanged. Importantly, inhibition of the proteasome inhibitor with MG132 did not recapitulate the increase in H2AK119ub following PSMD14 depletion **(Figure 4J)**, indicating that PSMD14 regulates H2A ubiquitination independently of its role in proteasomal degradation. Altogether, these results identify PSMD14 as a critical and direct regulator of H2AK119 ubiquitination in melanoma cells, revealing a previously unappreciated, proteasome-independent epigenetic function that may contribute to melanoma progression.

### PSMD14-dependent H2A deubiquitination activates pro-survival transcriptional programs in melanoma

Monoubiquitylation of histone H2A at lysine 119 (H2AK119ub) is an epigenetic mark associated with transcriptional repression and regulation of diverse biological processes, including cell proliferation and survival ^36^. To determine whether PSMD14 affects melanoma cell proliferation and survival through H2A-associated transcriptional programs, we used publicly available datasets on H2Aub-enriched genes ^37,38^. From these datasets, we selected 23 genes (hereafter referred to as H2Aub gene signature) involved in cellular pathways associated with melanoma progression **(Supplementary Table 2)**. Analysis of the TCGA melanoma dataset revealed a significant positive correlation between PSMD14 expression and the H2Aub gene signature (R=0.3743, p=3.329e-17) **(Figure 5A)**. In contrast, expression of the H2Aub-associated transcriptional program negatively correlated with RING1B expression (R=-0.325, p=3.97 e-13) **(Figure 5B)**, which encodes the Polycomb repressive complex 1 (PCR1) E3 ligase that catalyzes H2AK119 ubiquitination) ^37^. Consistent with these observations, endogenous expression of wild-type PSMD14, but not the catalytically inactive PSMD14 JAMM mut, resulted in increased expression of seven genes of the H2Aub gene signature (*MCL1*, *BCL2*, *TNFRSF19*, *PFKB3*, *SPP1*, *WNT10B*, and *HOXC4*) **(Figure 5C)**. Immunoblot analysis further confirmed that PSMD14 expression and enzymatic activity inversely correlated with global H2AK119ub levels and positively correlated with MCL1 and BCL2 protein levels **(Figure 5D, E)**. This relationship correlation was also found in 30 melanoma cell lines at the protein level (R=0.352, p=5.64 e-2) **(Supplementary Figure 3A)** and in a broad range of cancers at transcriptional level **(Supplementary Figure 3B)**. Conversely, PSMD14 depletion in A375 melanoma cells led to reduced expression of H2Aub-associated signature genes, including *MCL1*, *BCL2*, *TNFRSF19*, *PFKB3*, *SPP1*, *WNT10B*, and *HOXC4* **(Figure 5F)**, accompanied by increased H2AK119 ubiquitination and diminished MCL1 and BCL2 protein levels **(Figure 5G)**. ChIP-seq analysis further revealed enhance H2AK119ub occupancy across intronic and exonic regions of *MCL1* (chr1) and *BCL2* (chr 18) loci upon PSMD14 knockdown **(Supplementary Figure 3C)**, consistent with a repressed chromatin state and epigenetically mediated transcriptional silencing. MCL1 and BCL2 are major anti-apoptotic proteins ^39^. Therefore, we next examined the functional consequences of PSMD14-mediated H2A deubiquitination on melanoma cell survival. Time lapse monitoring of cell survival revealed that RING1B knock-down in melanoma cells rescued cell death induced by PSMD14 depletion **(Supplementary Figure 3D)**. Consistently, crystal violet assays showed that RING1B depletion partially restored cell viability in PSMD14-depleted A375 cells **(Figure 5H**). Immunoblot analyses confirmed that relative to PSMD14 knockdown alone, co-depletion of RING1B prevented H2AK119 ubiquitination, restored MCL1 and BCL2 expression, and reduced PARP cleavage, a marker of apoptotic cell death **(Figure 5I)**. Altogether, these findings indicate that PSMD14 and RING1B function antagonistically within the same pathway to regulate pro-survival gene expression through modulation of H2A ubiquitination.

**Figure 5.**
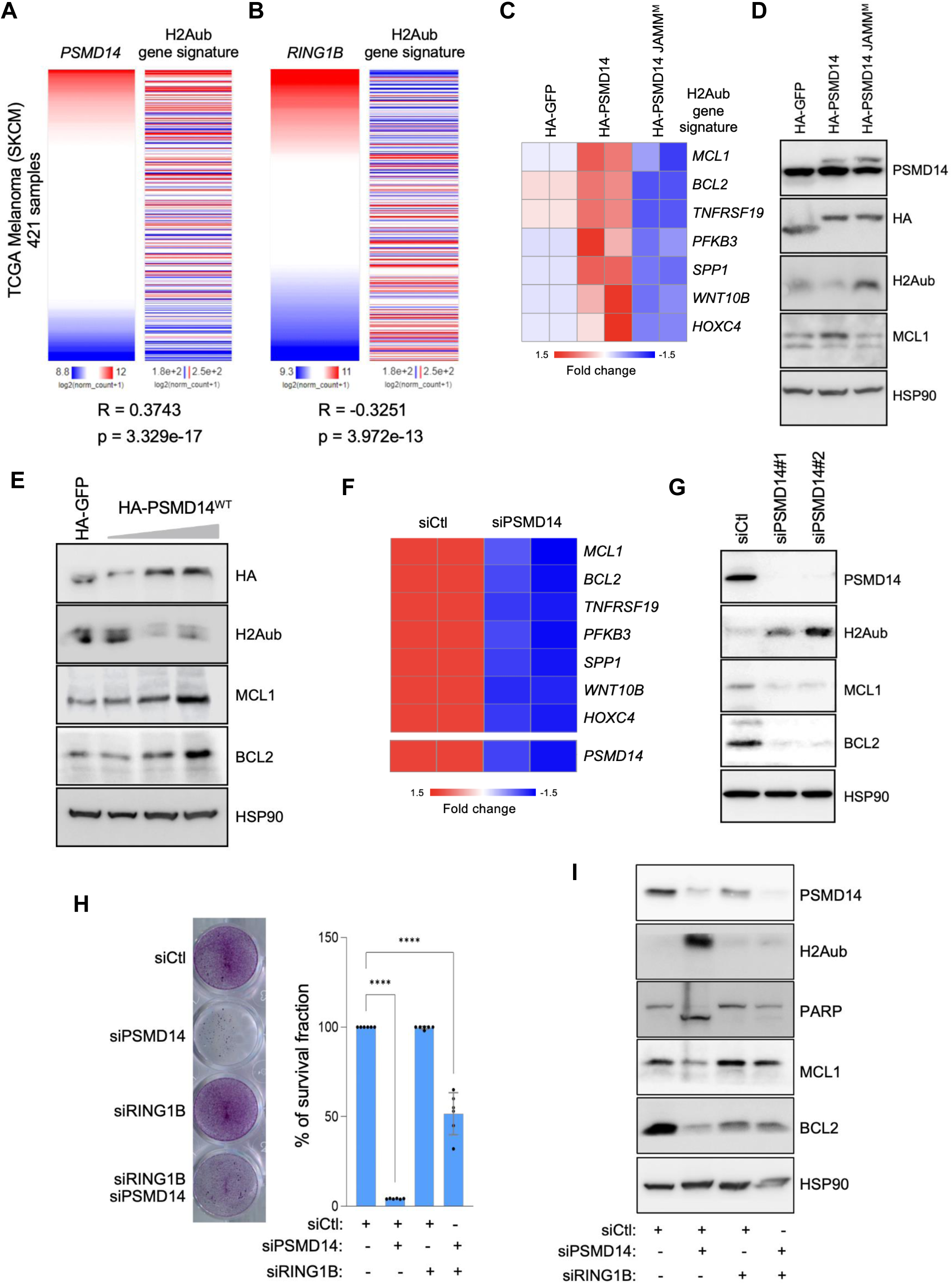
PSMD14 controls melanoma cell survival through H2A-associated transcriptional programs. **(A, B)** Correlation between the H2Aub gene signature (Supplementary Table X) and the expression of **(A)** *PSMD14* or **(B)** *RING1B* in the TCGA SKCM dataset (n = 421; UCSC Xena Browser). Correlations are assessed using Spearman’s rank-order test. **(C)** Heatmap showing mRNA expression of selected H2Aub target genes involved in apoptosis and metabolism in A375 melanoma cells stably expressing HA-PSMD14 or HA-PSMD14 JAMM^M^ compared to control cells (HA-GFP). **(D)** Western blot analysis showing the effects of increasing expression of HA-PSMD14 on H2Aub, MCL1 and BLC2 levels. **(E)** Western blot analysis of H2Aub and MCL1 levels in A375 cells expressing HA-PSMD14^WT^ or HA-PSMD14 JAMM^M^ compared to control cells (HA-GFP). **(F)** Heatmap showing expression changes in selected H2Aub target genes upon PSMD14 depletion in A375 cells. **(G)** Effect of PSMD14 knockdown (siPSMD14#1 and #2) in A375 cells on H2Aub, MCL1 and BLC2 levels compared to control (siCtl). HSP90, loading control. **(H)** Crystal violet staining (left) and quantification (right) of cell survival one week after knockdown of PSMD14 (siPSMD14) or RING1B (siRING1B) or the combined knockdown (siPSMD14/siRING1B). Bar graph shows cell survival relative to the control condition (siCtl). Data are mean ± SEM. Statistical significance was assessed using two-way ANOVA followed by Bonferroni’s multiple comparisons test. ****P ≤ 0.0001. **(I)** Western blot analysis of apoptotic markers (PARP, MCL1 and BCL2) and H2Aub levels following PSMD14 knockdown alone or in combination with RING1B depletion. HSP90, loading control.

### Targeting PSMD14/H2A axis enhances response to MAPK inhibition by preventing drug adaptation

We next examined the contribution of the PSMD14/H2A axis to melanoma cell responses to MAPK pathway inhibition. GSEA of eleven BRAF^V600^–mutant melanoma cell lines treated with the BRAF inhibitor Dabrafenib (GSE98314) revealed that, in addition to pathways linked to cell division and DNA replication, the Gene Ontology (GO) term protein deubiquitination (GO:0016579) was significantly associated with BRAF inhibition (NES=-2.030, p=3.01 e-16) **(Supplementary Figure 4A)**. Hierarchical clustering of genes associated with this pathway showed a marked downregulation of PSMD14 following MAPK inhibitor (MAPKi) treatment in BRAF-mutant melanoma cells **(Figure 6A)**. Consistently, PSMD14 mRNA levels were reduced upon treatment with BRAFi or MEKi alone, as well as with combined BRAFi/MEKi (vemurafenib/cobimetinib) therapy **(Supplementary Figure 4B and C)**. Immunoblot analysis further revealed that combined BRAFi/MEKi treatment inhibited ERK1/2 phosphorylation, reduced PSMD14 protein level, and increased H2AK119ub **(Figure 6B)**. The H2Aub gene signature was also significantly enriched in melanoma cells treated with Dabrafenib, either alone or in combination with the MEKi trametinib (GSE98314) (NES = -1.79, p= 8.1 e-4) **(Figure 6C)**. Of note, high levels of H2AK119ub were also observed in the NRAS-mutant cell line SBcl2 following increasing doses of MEKi Trametinib **(Supplementary Figure 4D)**, indicating that MAPK pathway inhibition promotes transcriptional repression through H2AK119 ubiquitination in both in BRAF- and NRAS-mutant melanomas.

**Figure 6.**
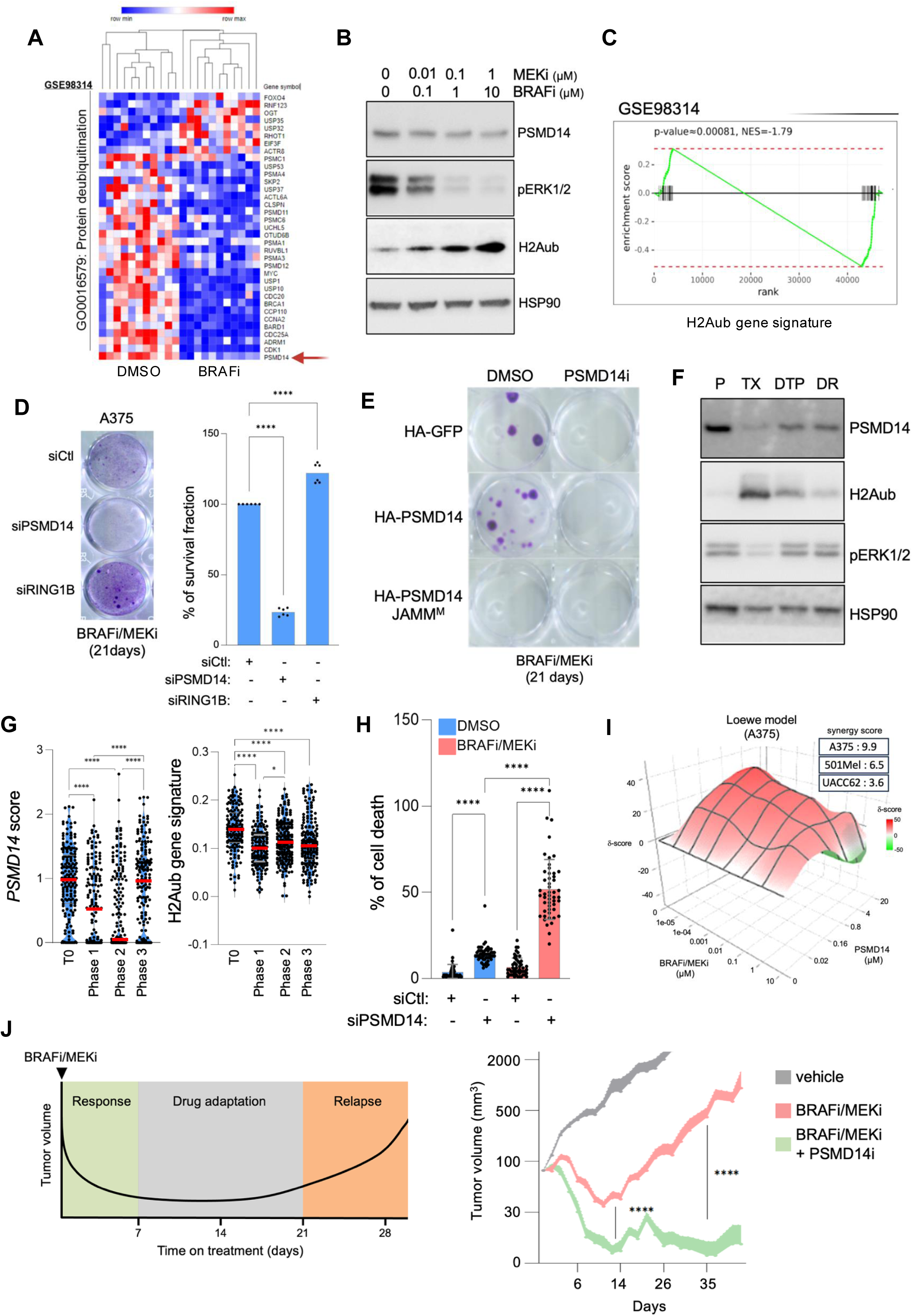
The PSMD14–H2Aub axis contributes to MAPK inhibitor response in melanoma. **(A)** Heatmap showing transcript levels of genes involved in protein deubiquitination biological process (GO:0016579) in BRAF-mutant melanoma cell lines treated with BRAF inhibitor dabrafenib with or without MEK inhibitor trametinib compared to DMSO (GSE98314). Each column represents one cell line. Red: high expression, blue: low expression. *PSMD14* is marked by a red arrow. HSP90, loading control. **(B)** Western blot analysis of PSMD14 and H2Aub levels in A375 cells treated with increasing doses and combinations of BRAFi and MEKi. HSP90, loading control. **(C)** Enrichment analysis of the H2Aub gene signature in BRAF-mutant melanoma cells exposed to BRAFi and/or MEKi (GSE98314). **(D)** Long-term crystal violet survival assay showing the effect of PSMD14 and RING1B depletion on residual cell survival after 21 days of BRAFi/MEKi treatment. Left, representative images of crystal violet cell staining. Right, bar graph showing the quantification of 2 independent experiments. Data are presented as mean ± SEM. Significance was determined with two-way ANOVA followed by Bonferroni’s multiple comparisons test. ****P ≤ 0.0001. **(E)** Long-term crystal violet survival assay showing the effect of HA-PSMD14^WT^ or HA-PSMD14 JAMM^M^ overexpression, in the presence or absence of PSMD14i (1 µM), on cell survival after 21 days of BRAFi/MEKi treatment. Cells were stained with crystal violet. A representative image of two independent experiments is shown. **(F)** Western blot analysis of PSMD14, H2Aub and P-ERK1/2 levels in A375 cells across stages of MAPKi treatment. P (untreated parental cells), TX (BRAFi/MEKi 24h), DTP (drug tolerant persister cells BRAFi/MEKi 21 days), DR (BRAFi/MEKi drug resistant cells). **(G)** *PSMD14* expression (left panel) and H2Aub gene signature (right panel) scores were extracted from single cell RNA sequencing of the MEL006 patient-derived xenograft model at drug-response phases (GSE116237) : pre-treatment (T0), 4 days of BRAFi/MEKi treatment (Phase 1), minimal residual disease (28 days on BRAFi/MEKi, Phase 2), and resistance phase (Phase 3). *P < 0.05. ****P < 0.0001. Two-way ANOVA.**(H)** Cell death analysis on A375 cells transfected with control siRNA (siCtl) or siPSMD14 and exposed or not to BRAFi/MEKi for 24 h. Cell death was monitored in real time using Incucyte® with Cytotox Red labelling. Data are mean ± SEM and statistical significance was assessed with two-way repeated-measures ANOVA. ****P ≤ 0.0001. **(I)** Synergy score determined according to the Loewe additivity model on A375 cells treated with the indicated combination of PSMD14i and BRAFi/MEKi. Scores >1 are characteristic of synergy above additivity and also visible in 501MEL and UACC62 BRAF-mutant melanoma cells (inset). **(J)** Schematic illustration of the A375 xenograft murine model of melanoma response to targeted therapies (left panel) and tumor volume changes in xenografts treated with BRAFi+MEKi (vemurafenib 30 mg/kg + cobimetinib 7 mg/kg, orally every 2 days) alone or combined with PSMD14i (8TQ, 15 mg/kg) (right panel). Vehicle-treated mice served as control. Group sizes were: Vehicle n = 10, BRAFi+MEKi n = 11, and BRAFi+MEKi+PSMD14i n = 8. Data represent mean ± SEM. \*\*\*\**P*<0.0001, two way repeated measures ANOVA followed by Bonferroni correction.

Since the emergence of drug-tolerant persister (DTP) cells is a critical intermediate step toward therapy resistance ^12,40^, we next examined whether the PSMD14-H2A axis contributes to melanoma cell persistence during MAPK treatment. A375 cells transfected with control, PSMD14 or RING1B siRNA were treated with BRAFi/MEKi for 21 days to allow DTP cell emergence. Compared with control cells, PSMD14 depletion drastically reduced the number of DTP cells, whereas RING1B knockdown significantly increased tolerance to BRAF/MEK inhibition **(Figure 6D)**. Conversely, overexpressing wild-type PSMD14, but not the inactive PSMD14 JAMM^M^ mutant, markedly increased drug persistence, an effect that was completely abrogated by pharmacological inhibition of PSMD14 with 8-TQ (PSMD14i) **(Figure 6E)**. We further examined PSMD14 expression and H2AK119ub levels in a model of BRAF-mutant A375 cells chronically exposed to BRAFi until resistance acquisition **(Figure 6F)**. Compared to drug-naïve cells (P), PSMD14 level was reduced during the early treatment-responsive phase (TX), consistent with the above observations **(Figure 6A, B)**. Importantly, PSMD14 expression progressively re-emerged during the DTP and fully drug-resistance (DR) phases, coinciding with decreased H2AK119ub levels and restored ERK signaling despite continued the presence of BRAFi/MEKi **(Figure 6F)**. *In silico* analysis of BRAF-mutant melanoma patient-derived xenografts (PDX) treated with BRAFi/MEKi until relapse (GSE116237) ^9^ further confirmed a dynamic correlation between PSMD14 level and the H2Aub gene signature throughout treatment **(Figure 6G)**. Given these observations, we examined whether disrupting the PSMD14-H2A axis could enhance responses to MAPK-targeted therapies. Cell survival analysis showed that PSMD14 depletion **(Figure 6H)** or pharmacological inhibition with 8-TQ **(Figure 6I)** synergized with BRAFi/MEKi to trigger melanoma cell death *in vitro*. Importantly, in a mouse model of transplanted melanoma, combining the PSMD14 inhibitor 8-TQ with BRAFi and MEKi markedly enhanced the early anti-tumor response and completely prevented tumor relapse, which typically occurs after three weeks of BRAFi/MEKi treatment **(Figure 6J)**. Altogether, these findings establish the PSMD14-H2A axis as a critical mediator of melanoma adaptation to MAPK inhibition and identify its targeting as an effective strategy to prevent therapy resistance *in vivo*.

## DISCUSSION

Growing evidence links DUBs to melanoma progression and therapeutic escap ^25^. However, the precise functions of many DUBs remain elusive ^24^. Here, we identified PSMD14 (RPN11, POH1) ^30–32^ as a critical regulator of melanoma cell survival, tumor development, therapeutic adaptation and drug resistance through a non-proteolytic regulation of histone H2A ubiquitination.

Our study builds on an unbiased genetic screen aimed at identifying DUBs essential for melanoma cell proliferation. Consistent with a recent study documenting the role of PSMD14 in melanoma growth ^33^, we demonstrate that catalytically active PSMD14 regulates the survival of a panel of melanoma cell lines harboring different mutational status of *BRAF*, *NRAS* and *NF1* through the prevention of proteotoxic stress and H2AX phosphorylation. Further supporting a major role of PSMD14 in melanoma biology, its pharmacological inhibition leads to DNA damage and apoptosis *in vivo* and impairs the development of BRAF and NRAS-mutant tumors in mice. While several studies have implicated PSMD14 as a proteasome-associated DUB that alters the stability of key players of DNA damage ^41^, and cell proliferation and survival pathways ^42,43^, the role of PSMD14 monodeubiquitinating activity in cancer development and chemoresistance is less understood. To gain insight on PSMD14 mechanism of action, we performed a proteomic analysis to investigate the PSMD14 interactome in melanoma. In addition to described components of the proteasome such as PSMD7 and PSMD12, several histone variants were found associated with PSMD14, including histone H2A members H2AJ, H2AFX and H2AY. H2A acts as a major epigenetic actor of transcriptional repression involved in cell fate and tumorigenesis ^44^. Previous work has established that monoubiquitylation of histone H2A by RING1 ubiquitin ligase within the Polycomb repressive complexes (*PRCs)* leads to transcriptionally silent and compacted chromatin ^45^. H2AK119 deubiquitinases, including BAP1 ^46^, MYSM1 ^47^ and USP22 ^48^ have been proposed as opposing this process. Adding to the list of DUBs sharing H2Aub as a common substrate, PSMD14 acts as a key H2AK119 deubiquitylase driving melanoma development and therapy resistance (this study) and myelomagenesis ^49^. In addition to H2A, our proteomic analysis of PSMD14 interactome in melanoma cells identified other histone variants (H2B, H3A, H4A) and proteins enriched in H2A nucleosomes including FACT complex subunits SPT16 and SSRP1 ^50^, consistent with the localization of PSMD14 with heterochromatin in yeast ^51^. These results further support the notion that PSMD14 is associated with protein complexes that display an affinity for H2A nucleosomes, and that these associations are important for H2A chromatin remodeling. Importantly, the H2A-directed activity of PSDM14 do not affect total level of H2A. Proteasome-associated PSMD14 is required to regulate poly-Ub conjugates generated at sites of DNA damage and regulates double-strand break response ^41^. Together with studies attributing an important role to nucleocytosolic translocation of the proteasome in cell survival under stress conditions ^52^, this raises the question of whether deubiquitination of H2AK119 by PSMD14 occurs independently of nuclear proteasomes.

Molecular therapy targeting the oncogenic BRAF pathway has improved melanoma patient survival, but resistance linked to incomplete tumor response and residual disease inevitably develops ^6^. Multiple genetic and non-genetic mechanisms of melanoma resistance to TT have been described ^4,12^. However, the epigenetic regulation of drug resistance remains poorly understood. In addition to its role in melanoma cell survival, we show here that the PSDM14/H2A axis plays an important role in cell adaptation and resistance to targeted therapies. Whereas PSMD14 inhibition or depletion enhances the anti-melanoma action of BRAF/MEK inhibitors, overexpression of PSMD14, but not of a catalytically inactive JAMM mutant, improved cancer cell survival and the ability to develop drug resistance. On the other hand, RING1B depletion enhances melanoma cell persistence in the presence of BRAF/MEK inhibitors. Deubiquitination of H2AK119 by PSMD14 was associated with a transcriptional program (the ‘’H2Aub gene signature’’) that includes genes encoding for cell survival proteins, such as the anti-apoptotic proteins BCL2 and MCL1 ^53^. Conversely, targeting PSMD14 increases H2AK119ub-dependent program associated with increased DNA damage, reduced MCL1 and BCL2 levels, cell proliferation arrest and death, thereby enhancing response to MAPK pathway inhibition and reducing tumor relapse. The clinical relevance of our findings is strongly supported by the fact that the recently developed MCL1 inhibitor AZD5991 delays acquired resistance to BRAF/MEK inhibitors, suggesting that MCL1 is a driver of adaptive survival in BRAF-mutated melanoma treated with targeted therapies ^54^. Early treatment of melanoma cells with BRAFi/MEKi also promotes H2AK119 ubiquitination by triggering PSMD14 downregulation, while PSMD14 levels and H2Aub gene signature are progressively restored in the drug-tolerant and drug-resistant states. Combining a PSMD14 inhibitor with BRAF/MEK inhibitors is likely to abrogate this tolerant phase, thereby overcoming melanoma drug resistance. These observations strongly support the notion that tuning H2A ubiquitination levels through PSMD14 (and possibly other H2A DUBs) in a timely manner is essential to maintain chromatin plasticity during the process of adaptation to therapeutic stress that precedes drug resistance ^12,55^. In this context, it would be important to investigate how PSMD14 expression is restored during the tolerant phase in the presence of targeted therapies. In addition, it remains unclear whether TT-induced, H2A-mediated epigenetic reprogramming also entails additional histone modifications—such as methylation—that have been linked to BRAF inhibition ^56^ and implicated in chromatin-dependent regulation of the drug-tolerant state ^55^. Interestingly, haploinsufficiency of the histone deacetylase SIRT6 induces BRAF-mutant melanoma cell resistance to TT via H3K56 acetylation at the IGFBP2 locus and increased IGF-1R/AKT signaling ^57^, suggesting that epigenetic mechanisms of drug resistance are multiple.

Collectively, our study not only highlights the crucial role of the PSMD14/H2A axis for melanoma cell survival and tumor development but also identifies a novel epigenetic mechanism of drug resistance. Targeting PSMD14 in combination with standard treatment for BRAF-mutated melanoma may thus represent an effective strategy for treating aggressive melanoma and countering therapeutic resistance.

## MATERIALS AND METHODS

### Cell culture and reagents

MeWo, A375, WM793 and UACC62 human melanoma cell lines and HEK293T cells were from the American Type Culture Collection (ATCC). WM164 cells were purchased from Rockland. WM9 and Sbcl2 cells were provided by M. Herlyn. M229, M238 and M249 melanoma cell lines and their corresponding BRAF inhibitor-resistant sublines were obtained from R. Lo ^10^. The 501Mel melanoma cell line was a gift from R. Halaban. The A375DDR double-resistant variant was generated from parental A375 cells by repeated exposure to the BRAF inhibitor vemurafenib and the MEK inhibitor trametinib for 5 to 6 months until stable resistance was established. Murine melanoma primary cell lines YUMM1.7 (*Braf*V600E/*Pten* null/*Cdkn2a* null) ^58^ and MaNRAS (*NRAS*Q61K) ^59^ were obtained from M. Bosenberg (Yale School of Medicine, USA) and L. Larue (Institut Curie, France), respectively. Short-term patient-derived melanoma cells MM099 and MM029 were kindly provided by J.-C. Marine (VIB, Belgium). Human melanoma cell lines and HEK293T cells were cultured in Dulbecco’s modified Eagle’s medium (DMEM) or RPMI-1640, as appropriate, supplemented with 7% fetal bovine serum (FBS; GE Healthcare HyClone), 50 U/mL penicillin, and 50 µg/mL streptomycin. YUMM1.7 cells were maintained in Opti-MEM medium (Thermo Fisher Scientific) supplemented with 3% FBS and penicillin/streptomycin. MaNRAS cells were cultured in Ham’s F-12 medium supplemented with 10% fetal calf serum (FCS), penicillin/streptomycin, and 100 nM phorbol 12-myristate 13-acetate (PMA). All cell lines were cultured at 37°C in a humidified atmosphere containing 5% CO₂ and were routinely tested for mycoplasma contamination by PCR. Lipofectamine RNAiMAX and Opti-MEM medium were purchased from Invitrogen. Vemurafenib (PLX4032) and trametinib (GSK1120212) were obtained from Selleckchem. The PSMD14 inhibitor 8-(tosylamino)quinoline (8-TQ) ^32^ was purchased from Merck.

### Plasmids, siRNA and transfection

Flag-HA-GFP (Addgene plasmid #22612; http://n2t.net/addgene:22612; RRID:Addgene_22612) and Flag-HA-PSMD14 (Addgene plasmid #22557; http://n2t.net/addgene:22557; RRID:Addgene_22557) were a gift from Wade Harper (Sowa et al., 2009). The PSMD14-JAMM mutant was generated by substituting histidines at residues 113 and 115 with alanines ^31^ using the QuikChange site-directed mutagenesis kit (Agilent). The primers used for PSMD14-JAMM plasmid generation were as follows: forward, 5’-GGCCTCTACCAACAACCATACGATCACGGACCGAAACCAACAACCG-3’; reverse, 5’-GGAGATGGTTGTTTTGGTATGCTAGTGCCCCTGGCTTTGGTTGGC-3’. All constructs were verified by Sanger sequencing. The pCDNA3.1-Flag-H2A plasmid (Addgene plasmid #63560; http://n2t.net/addgene:63560 ; RRID:Addgene_63560) was a gift from Titia Sixma. Transient plasmid transfections were performed in HEK293T and A375 cells using Lipofectamine 2000 (Thermo Fisher Scientific), according to the manufacturer’s instructions. Stable A375-GFP, A375-PSMD14, and A375-PSMD14-JAMM cell lines were generated by transfecting A375 cells with the indicated plasmids, followed by puromycin selection (InvivoGen). All siRNA transfections were performed using Lipofectamine RNAiMAX (Thermo Fisher Scientific) according to the manufacturer’s recommendations. For the siRNA screen, a siGENOME® SMARTpool® siRNA Library Human Deubiquitinating Enzymes (Catalog #G-104705-01; Dharmacon, Horizon Discovery) was used. Briefly, 501Mel cells (1 × 10⁵ cells per well) were seeded in 24-well plates and transfected 24 h later with 50 nM non-targeting SMARTpool® siRNA or individual DUB-targeting SMARTpool® siRNAs. Cell proliferation was monitored by live-cell imaging using the IncuCyte ZOOM™ system (Essen BioScience) for 72 h. For targeted RNA interference experiments, cells were seeded on culture dishes and transfected 24 h later with 50 nM control siRNA (Universal Negative Control #1, SIC001), PSMD14 siRNAs (SASI_Hs02_00340316, SASI_Hs01_00024446, SASI_Hs01_00024447), or RNF2 siRNA (SASI_Hs01_00213463) (Merck), diluted in serum-free medium containing Lipofectamine RNAiMAX. Cells were incubated at 37°C in a humidified atmosphere containing 5% CO₂ for the indicated times post-transfection.

### Immunofluorescence staining

For phospho-histone H2AX foci detection, cells grown on glass coverslips were fixed with 4% paraformaldehyde and permeabilized in buffer containing 50 mM Tris-HCl (pH 7.4), 150 mM NaCl, and 0.3% Triton X-100 for 1 h at 4°C. Cells were then blocked in 50 mM Tris-HCl (pH 7.4), 150 mM NaCl, 0.03% Triton X-100, and 1% bovine serum albumin (BSA) for 1 h at 37°C. Cells were incubated overnight at 4°C with an anti-phospho-histone H2AX (Ser139) antibody (1:1000; Merck Millipore, Cat# 05-636). After washing, cells were incubated for 1 h at room temperature with Alexa Fluor™ Texas Red–conjugated secondary antibodies and counterstained with 10 µg/mL DAPI (Merck, Cat# D9542). After washing, coverslips were mounted using ProLong™ Gold Antifade Mountant (Thermo Fisher Scientific). Immunofluorescence images were acquired using a Zeiss confocal microscope equipped with a 40× objective. Image acquisition was performed using the Zeiss LSM Browser software. Quantitative image analysis was carried out using ImageJ software (NIH). Cells were considered phospho-H2A.X–positive when nuclei exhibited at least a fivefold increase in γH2AX foci compared to control cells.

### Immunohistochemistry

Paraffin-embedded tumor tissues were sectioned at 4 µm thickness. Tissue sections were dewaxed, rehydrated, and subjected to heat-induced antigen retrieval using the PT Module Lab Vision™ (Thermo Fisher Scientific), according to the manufacturer’s instructions. Sections were then incubated with primary antibodies diluted in 1% (wt/vol) bovine serum albumin (BSA), followed by standard avidin–biotin–peroxidase complex (ABC) staining using the automated Myreva SS30 staining system (Microm Microtech, France). Primary antibodies used were as follows: phospho-histone H2A.X (Ser139; γH2AX; Merck Millipore, Cat# 05-636), Ki67 (Abcam, Cat# ab16667), and cleaved caspase-3 (Asp175; Cell Signaling Technology, Cat# 9661).

### Cell proliferation and long-term survival assays

Cell proliferation and survival were assessed using crystal violet staining at both short- and long-term time points. For short-term endpoint proliferation assays (≤72 h), cells were fixed with 3% paraformaldehyde (PFA) for 20 min, washed with phosphate-buffered saline (PBS), and stained with 0.4% crystal violet dissolved in 20% ethanol for 30 min at room temperature. Excess dye was removed by extensive washing with water, and bound crystal violet was solubilized using 10% acetic acid. Absorbance was measured at 595 nm using a microplate reader. Relative cell number was expressed as a percentage of the absorbance measured in control conditions.

For long-term survival assays (14–21 days), cells were treated as indicated and maintained in culture over extended periods, with medium and treatments renewed every 3–4 days. At the end of the treatment period, cells remaining adherent to the culture dish were fixed and stained using the same crystal violet protocol. Representative images were acquired, and residual cell survival was quantified by measuring crystal violet staining intensity using ImageJ software. Data were expressed relative to control conditions. These long-term assays reflect cumulative cell survival rather than clonal outgrowth.

Alternatively, cell proliferation was monitored in real time using the IncuCyte ZOOM™ live-cell imaging system (Essen BioScience). Phase-contrast images were acquired every hour for up to 72 h, and cell growth curves were generated using the IncuCyte software based on cell confluence. Cell proliferation at 72 h was expressed relative to control conditions.

### Cell viability assays

After the indicated treatment times, cell viability was assessed by flow cytometry following staining with Annexin V–FITC and propidium iodide (PI) (Thermo Fisher Scientific). Flow cytometry data were acquired using a MACSQuant® flow cytometer (Miltenyi Biotec) and analyzed with MACSQuantify™ software (Miltenyi Biotec). Gating strategies were applied to exclude debris and doublets prior to quantification of Annexin V–FITC- and DAPI-positive cell populations. The percentages of apoptotic and dead cells were determined based on Annexin V–FITC and DAPI staining. For IC₅₀ determination, melanoma cells were treated with increasing concentrations of the PSMD14 inhibitor 8-TQ, and cell viability was quantified by Annexin V–FITC/PI staining 72 h after treatment. IC₅₀ values were calculated using GraphPad Prism software (GraphPad Software) by nonlinear regression analysis of dose–response curves using the log[inhibitor] versus response (variable slope) model, with default Prism parameters.

Real-time analysis of cell death was performed using the IncuCyte ZOOM™ live-cell imaging system by monitoring Cytotox Red fluorescence as a readout of loss of membrane integrity, according to the manufacturer’s instructions.

### Synergistic determination of drug combination effects on cell viability

To assess potential synergistic effects of inhibitor combinations, A375 cells were seeded at a density of 0.5 × 10⁴ cells per well in 48-well plates (Corning) and treated with either single inhibitors or their combinations at the indicated concentrations, in technical triplicates. Dilution series were predetermined based on the IC₅₀ values of each inhibitor. Cell viability was assessed after 48 h of treatment using the IncuCyte® real-time live-cell analysis system, based on cell confluence measurements. Synergy scores for drug combinations were calculated using the Loewe reference model implemented in SynergyFinder 3.0 (https://synergyfinder.fimm.fi).

### mRNA preparation and real-time qPCR

Total RNA was extracted from cell samples using the RNeasy Micro Kit (Qiagen, Cat# 74004) according to the manufacturer’s instructions. Two micrograms of total RNA were reverse transcribed into cDNA using the GoScript™ Reverse Transcription System (Promega, Cat# A5001). Quantitative real-time PCR was performed using Platinum SYBR Green qPCR SuperMix (Thermo Fisher Scientific, Cat# 4309155) on a StepOne™ Real-Time PCR System (Applied Biosystems). RSP14 gene expression was used for normalization, and relative gene expression levels were calculated using the comparative ΔΔCt method using StepOne software v2.1. Primer sequences were designed using the PrimerBank database (https://pga.mgh.harvard.edu/primerbank/). Heatmap visualization of differential gene expression was performed using MeV software (http://mev.tm4.org/).

### Chromatin immunoprecipitation followed by sequencing (ChIP-seq)

ChIP-seq experiments were performed by Active Motif (Carlsbad, CA). Briefly, chromatin was prepared from formaldehyde-fixed cells by cell lysis, followed by washing and sonication using the PIXUL® Multi-Sample Sonicator (Active Motif, Cat# 53130) to shear DNA to an average fragment size of 200–1000 bp. To assess chromatin yield, an aliquot of sheared chromatin was crosslinked at 65°C, treated with RNase and proteinase K, and DNA was purified using SPRI beads (Beckman Coulter). DNA concentration was measured using a Qubit Fluorometer (Thermo Fisher Scientific), and total chromatin yield was extrapolated from the initial chromatin volume. For chromatin immunoprecipitation, chromatin aliquots were precleared with protein G agarose beads (Invitrogen). Immunoprecipitations were performed using an antibody specific for ubiquitinated histone H2A (H2AK119ub; Cell Signaling Technology, Cat# 8240; 4 µg per condition). Following extensive washing, immune complexes were eluted using SDS-containing buffer, treated with RNase and proteinase K, and de-crosslinked by overnight incubation at 65°C. ChIP DNA was purified by phenol–chloroform extraction followed by ethanol precipitation. ChIP DNA libraries were prepared using either the PrepX DNA Library Kit (Takara Bio) on the Apollo™ automation platform or the NEB DNA Library Prep Kit (New England Biolabs), according to the manufacturers’ instructions.

Libraries were sequenced on an Illumina platform, and raw sequencing data were processed using standard ChIP-seq analysis pipelines. For visualization of ChIP-seq signal enrichment and peak distribution across genomic loci, normalized coverage tracks were generated and visualized using the Integrative Genomics Viewer (IGV, Broad Institute). ChIP-seq signal profiles at target gene loci, including MCL1 and BCL2, were displayed as genome browser tracks to compare H2AK119ub occupancy between experimental conditions.

### Western blot analysis

Whole-cell lysates were prepared using ice-cold RIPA buffer supplemented with protease and phosphatase inhibitor cocktails (Pierce™, Thermo Fisher Scientific, Cat# 78440 and Cat# 78420). Lysates were clarified by centrifugation at 14,000 rpm for 20 min at 4°C. Protein concentrations were determined using standard procedures, and 30 µg of total protein were separated by SDS-PAGE and transferred onto PVDF membranes (Amersham™ Hybond™ PVDF, Merck, Cat# GE10600023). Membranes were blocked and incubated with primary antibodies overnight at 4°C in saturation buffer (5% BSA, 1 mM EDTA, 500 mM NaCl, 10 mM Tris-HCl, pH 7.4). Primary antibodies used were as follows: PSMD14 (Elabscience, Cat# E-AB-63456), p21^Waf1/Cip1 (clone 12D1; Cell Signaling Technology, Cat# 2947), HSP90 (Santa Cruz Biotechnology, Cat# sc-13119), PARP (Cell Signaling Technology, Cat# 9542S), Cleaved Caspase-3 (Asp175; Cell Signaling Technology, Cat# 9661), γH2AX (Cell Signaling Technology, Cat# 9718S), H2Aub (Lys119) (clone D27C4; Cell Signaling Technology, Cat# 8240S), ubiquitinylated proteins (Merck Millipore, Cat# 04-263), Histone H2A (Cell Signaling Technology, Cat# 12349S), HA tag (Sigma-Aldrich, Cat# H9658), PSMD7 (Santa Cruz Biotechnology, Cat# sc-390705), PSMD12 (Santa Cruz Biotechnology, Cat# sc-398279), MCL-1 (Cell Signaling Technology, Cat# 94296), BCL-2 (Cell Signaling Technology, Cat# 15071), and phospho-ERK1/2 (Thr202/Tyr204; Cell Signaling Technology, Cat# 9101). After washing with TBS-T buffer (10 mM Tris-HCl, pH 7.5, 500 mM NaCl, 0.1% Tween-20), membranes were incubated for 1 h at room temperature with horseradish peroxidase (HRP)-conjugated secondary antibodies (Cell Signaling Technology). Protein bands were detected by enhanced chemiluminescence using ECL reagents (Amersham™ ECL™, Cytiva). Western blots shown are representative of at least three independent experiments.

### Immunopurification and sample preparation for mass spectrometry

Whole-cell lysates from A375 cells stably expressing Flag-HA-PSMD14 or Flag-HA-GFP were prepared using ice-cold RIPA buffer supplemented with protease and phosphatase inhibitor cocktails (Pierce™, Thermo Fisher Scientific, Cat# 78440 and Cat# 78420). Lysates were generated from approximately 5 × 10⁷ cells per condition. Anti-HA immunoaffinity columns were prepared using anti-HA agarose beads (Sigma-Aldrich/Merck, Cat# A2095) according to the manufacturer’s instructions. Cleared lysates were applied to the anti-HA agarose and incubated overnight at 4°C with gentle rotation. Immunoprecipitated complexes were washed three times with 10 mM Tris-HCl, pH 7.5, 500 mM NaCl, 0.05% Tween-20 and eluted in Laemmli sample buffer (Thermo Fisher Scientific, Cat# 39000). As a quality control, a fraction of the immunopurified material was analyzed by immunoblotting using an anti-HA tag antibody (Cell Signaling Technology, Cat# 3724).

### Mass Spectrometry analysis, protein identification and quantification

The immunoprecipitated samples were loaded on NuPAGE™ 4–12% Bis–tris acrylamide gels according to the manufacturer’s instructions (Invitrogen, Life Technologies). Running of samples was stopped as soon as proteins stacked as a single band. Protein containing bands were stained with Thermo Scientific Imperial Blue, cut from the gel, and following reduction and iodoacetamide alkylation, digested with high sequencing grade trypsin (Promega, Madison, WI, USA). Extracted peptides were concentrated before mass spectrometry analysis. Samples were reconstituted with 0.1% trifluoroacetic acid in 2% acetonitrile and analyzed by liquid chromatography (LC)-tandem MS (MS/MS) using a Q Exactive Plus Hybrid Quadrupole-Orbitrap online with a nanoLC Ultimate 3000 chromatography system (Thermo Fisher Scientific™, San Jose, CA). For each biological sample (prepared in triplicate), 5 microliters corresponding to 20 % of digested sample were injected in triplicate on the system. After pre-concentration and washing of the sample on a Acclaim PepMap 100 column (C18, 2 cm × 100 μm i.d. 100 A pore size, 5 μm particle size), peptides were separated on a LC EASY-Spray column (C18, 50 cm × 75 μm i.d., 100 A, 2 µm, 100A particle size) at a flow rate of 300 nL/min with a two steps linear gradient (2-22% acetonitrile/H20; 0.1 % formic acid for 100 min and 22-32% acetonitrile/H20; 0.1 % formic acid for 20 min). For peptides ionization in the EASYSpray source, spray voltage was set at 1.9 kV and the capillary temperature at 250 °C. All samples were measured in a data dependent acquisition mode. Each run was preceded by a blank MS run in order to monitor system background. The peptide masses were measured in a survey full scan (scan range 375-1500 m/z, with 70 K FWHM resolution at m/z=400, target AGC value of 3.00×106 and maximum injection time of 100 ms). Following the high-resolution full scan in the Orbitrap, the 10 most intense data-dependent precursor ions were successively fragmented in HCD cell and measured in Orbitrap (normalized collision energy of 25 %, activation time of 10 ms, target AGC value of 1.00×105, intensity threshold 1.00×104 maximum injection time 100 ms, isolation window 2 m/z, 17.5 K FWHM resolution, scan range 200 to 2000 m/z). Dynamic exclusion was implemented with a repeat count of 1 and exclusion duration of 20 s.

For protein identification and quantification, relative intensity-based label-free quantification (LFQ) was processed using the MaxLFQ algorithm from the freely available MaxQuant computational proteomics platform, version 1.6.3.4. Analysis was done on three biological replicates, each injected three times on mass spectrometers. The acquired raw LC Orbitrap MS data were first processed using the integrated Andromeda search engine. Spectra were searched against the against the human protein proteome of the swissprot database (20,368 entries, extracted from Uniprot on november 2019). The false discovery rate (FDR) at the peptide and protein levels were set to 1% and determined by searching a reverse database. For protein grouping, all proteins that could not be distinguished based on their identified peptides were assembled into a single entry according to the MaxQuant rules. The statistical analysis was done with Perseus program (version 1.6.15) from the MaxQuant environment (www.maxquant.org). Quantifiable proteins were defined as those detected in above 70% of samples in one condition or more. Protein LFQ normalized intensities were base 2 logarithmized to obtain a normal distribution. Missing values were replaced using data imputation by randomly selecting from a normal distribution centered on the lower edge of the intensity values that simulates signals of low abundant proteins using default parameters (a downshift of 1.8 standard deviation and a width of 0.3 of the original distribution). To determine whether a given detected protein was specifically differential in HA-PSMD14 versus HA-GFP immunopreciptation, a two-sample *t*-test was done using permutation-based FDR-controlled at 0.5 % and employing 250 permutations. The *p-*value was adjusted using a scaling factor s0 with a value of 1.

### *In vivo* experimentation

All mouse experiments were conducted in accordance with institutional guidelines and approved by the local ethics committee for animal experimentation (CIEPAL-Azur; agreement NCE/2018-509), in compliance with national and european regulations. For syngeneic melanoma models, 5 × 10⁵ YUMM1.7 or MaNRAS murine melanoma cells were subcutaneously injected into the flanks of 6-week-old female C57BL/6J mice (Envigo). When tumors became palpable (approximately 0.05–0.1 cm³), mice were treated intraperitoneally every other day with vehicle or the PSMD14 inhibitor (8-TQ, 15 mg/kg), formulated in a 90:9:1 (v/v/v) mixture of Labrafil, dimethylacetamide, and Tween 80. For xenograft melanoma models, 2 × 10⁶ A375 human melanoma cells were subcutaneously inoculated into 6-week-old immunodeficient athymic Foxn1nu/nu mice (Janvier Labs). Once tumors reached a volume of approximately 0.1 cm³, mice received oral administration of vemurafenib (30 mg/kg) and cobimetinib (7 mg/kg) every other day, or matched vehicle control. After the initial tumor response phase, animals were randomized into two treatment groups: one group continued to receive vemurafenib and cobimetinib alone, while the second group received the same combination together with daily intraperitoneal injections of 8-TQ (10 mg/kg). Tumor growth was monitored by caliper measurements, and tumor volume was calculated using the formula: volume = (length × width²)/2. At the experimental endpoint, or when tumor volume reached 1 cm³, mice were euthanized according to institutional guidelines, and tumors were dissected, weighed, fixed, and paraffin-embedded for subsequent immunohistochemical analyses.

### In silico analyses

Melanoma datasets from The Cancer Genome Atlas (TCGA, Skin Cutaneous Melanoma [SKCM]) were analyzed using the GEPIA platform (http://gepia.cancer-pku.cn/) to compare PSMD14 expression levels between normal skin and primary melanoma samples. Publicly available Gene Expression Omnibus (GEO) datasets were used to assess PSMD14 expression across melanoma progression, including the GSE3189 dataset. Normalized expression data were analyzed using GraphPad Prism v5.0b (GraphPad Software). Overall survival analyses were performed using TCGA melanoma patient data retrieved from the SurvExpress platform (http://bioinformatica.mty.itesm.mx:8080/Biomatec/SurvivaX.jsp). Patients were stratified into high- and low-risk groups based on the prognostic index (PI), calculated using Cox proportional hazards regression, with equal numbers of samples in each group after ordering by PI values. Gene Set Enrichment Analysis (GSEA) was performed on TCGA Skin Cutaneous Melanoma Firehose Legacy datasets (n = 472 patients) to compare PSMD14^high^ versus PSMD14^low^ melanoma samples using the GSEA software (http://software.broadinstitute.org/gsea/index.jsp). A gene signature corresponding to genes transcriptionally silenced by histone H2A ubiquitination (H2Aub gene signature; Supplementary Table XX) was generated from publicly available datasets ^37,38^. Expression of genes associated with the protein deubiquitination Gene Ontology term (GO:0016579), together with the H2Aub signature, was evaluated in BRAF-mutant melanoma cell lines treated with BRAF and MEK inhibitors (dabrafenib and trametinib) using the GSE98314 dataset. Normalized enrichment scores (NES) and heatmap visualizations were generated using Phantasus (https://ctlab.itmo.ru/phantasus/). Gene dependency scores derived from genome-scale CRISPR-Cas9 (Avana) screens were obtained from the DepMap portal (https://depmap.org/) using CERES-corrected gene effect values across melanoma and pan-cancer cell lines.

## Statistical analysis

Unless otherwise stated, all experiments were performed at least three independent times, and representative data or images are shown. Statistical analyses were performed using GraphPad Prism software (GraphPad Software). Data are presented as mean ± SEM. For comparisons between two groups, statistical significance was assessed using unpaired two-tailed Student’s t-tests, or Mann–Whitney tests when data did not follow a normal distribution. Comparisons involving more than two groups were analyzed using one-way ANOVA or two-way ANOVA, as appropriate, followed by Bonferroni’s multiple comparisons test. Longitudinal data, including tumor growth and real-time cell proliferation assays, were analyzed using two-way repeated-measures ANOVA. For proteomics analyses, differential protein enrichment was assessed using a two-sample t-test with permutation-based false discovery rate (FDR) correction. P values < 0.05 were considered statistically significant and are indicated as follows: *P < 0.05, **P < 0.01, ***P < 0.001, \*\*\*\**P*<0.0001.

## Data availability

The mass spectrometry proteomics data have been deposited on the ProteomeXchange Consortium of proteomics resource (http://www.proteomexchange.org) with the dataset identifier PXD073113.

## Acknowledgments

We thank J.C. Marine for the short-term cultured melanoma cells, M. Irondelle from the C3M imaging facility and members of the C3M animal facility. This work was supported by funds from Inserm, Société Française de Dermatologie, Fondation ARC pour la recherche sur le cancer and Ligue contre le cancer. The financial contribution of the Conseil général 06, Canceropôle Provence Alpes Côte d’Azur and Région Provence Alpes Côte d’Azur to the C3M is also acknowledged. The Marseille Proteomic facility (MaP; http://map.univmed.fr/) is supported by IBiSA (Infrastructures Biologie Santé et Agronomie), Canceropôle PACA, Région PACA and Institut Paoli-Calmettes, Fonds Européen de Développement Regional (FEDER) and Plan Cancer. P.B. is a recipient of a doctoral fellowship from Fondation ARC pour la recherche sur le cancer. M.K. is a recipient of a doctoral fellowship from Ligue Nationale Contre le Cancer.

## Author contributions

M.D. and M.O. designed the study. M.O. performed the experiments and analyzed the data with the help of P.B., M.K., S.D., L.L. and F.L. R.D. performed the initial genetic screen. S.A. performed the mass spectrometry analysis. V.D. and L. L. provided the NRAS animal model and comments on the manuscript. M.D. and M.O. wrote the original draft. M.D. supervised the study and edited the final version of the manuscript with the help of L.L. and S.T.-D.

## Supplementary Figure Legends

**Supplementary Figure 1.**
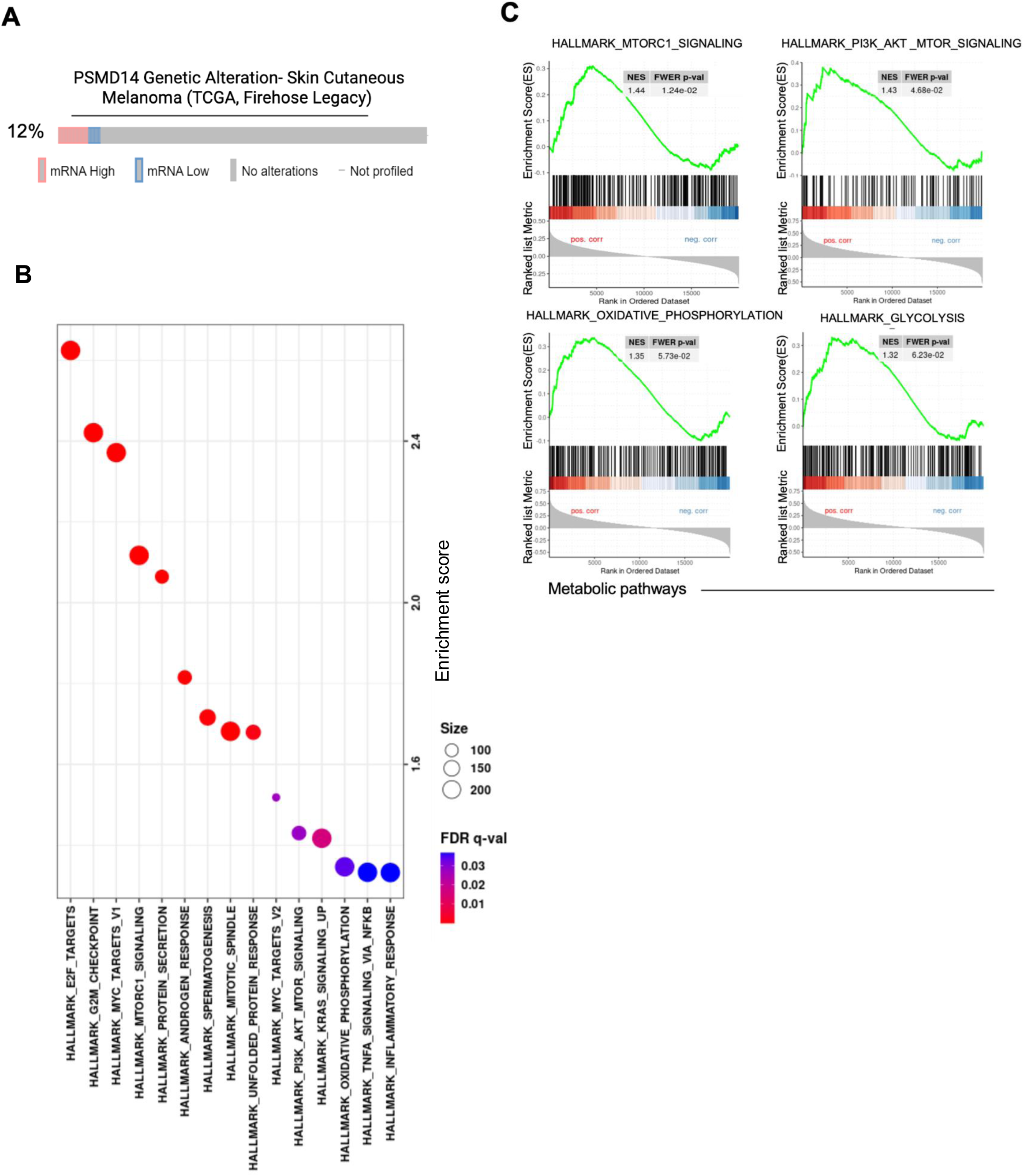
**(A)** Proportion of *PSMD14* alterations in 471 SKCM patient samples from the cBioPortal database. **(B)** Hallmark enrichment and GSEA analyses associated with *PSMD14* expression. **(C)** TCGA skin melanoma data showing a significant correlation between PSMD14 expression and melanoma aggressiveness–related hallmarks.

**Supplementary Figure 2.**
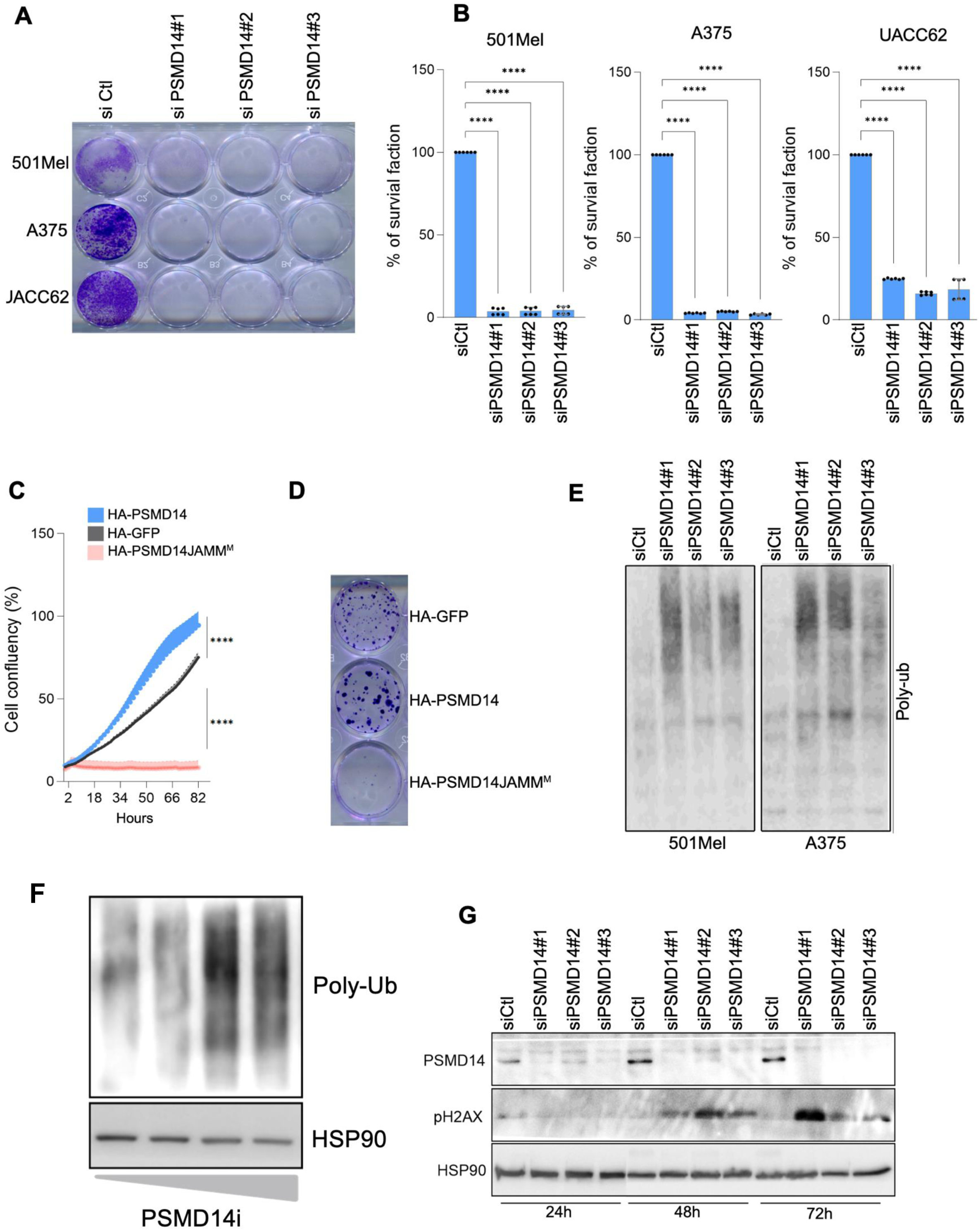
**(A)** Representative crystal violet–stained images showing residual adherent cells following PSMD14 depletion (siPSMD14 #1, #2, and #3) versus control siRNA (siCtl) in UACC62, A375, and 501Mel cells, followed by crystal violet staining and quantification. **(B)** Quantification of long-term crystal violet survival assays shown in (A). Bar graphs show fold-change of cell survival in siPSMD14 treated cells relative to control (siCtl). Data are the mean ± SEM (n=3). \*\*\*\**P*<0.0001, two-way ANOVA. **(C)** Time-lapse analysis of cell confluency in A375 cells expressing wild-type (WT) or mutant (JAMM^M^) HA-PSMD14, compared with HA-GFP controls. Statistical significance for longitudinal proliferation curves was assessed using a two-way repeated-measures ANOVA . \*\*\*\**P*<0.0001. **(D)** Long-term crystal violet survival assay showing the effect of HA-GFP, HA-PSMD14^WT^ and HA-PSMD14 JAMM^M^ overexpression on A375 cell survival. Cells were stained with crystal violet after 2 weeks. A representative image of two independent experiments is shown**. (E)** Levels of poly-ubiquitinated proteins (poly-Ub) in 501Mel and A375 cells depleted of PSMD14 (siPSMD14 #1, #2, and #3), assessed by Western blot. **(F)** Levels of poly-ubiquitinated proteins (poly-Ub) in A375 cell lysates treated with increasing concentrations of PSMD14i 8-TQ, assessed by Western blot. HSP90, loading control. **(G)** Western blot analysis of γH2AX levels in PSMD14-depleted cells at 24 h, 48 h, and 72 h. HSP90, loading control.

**Supplementary Figure 3.**
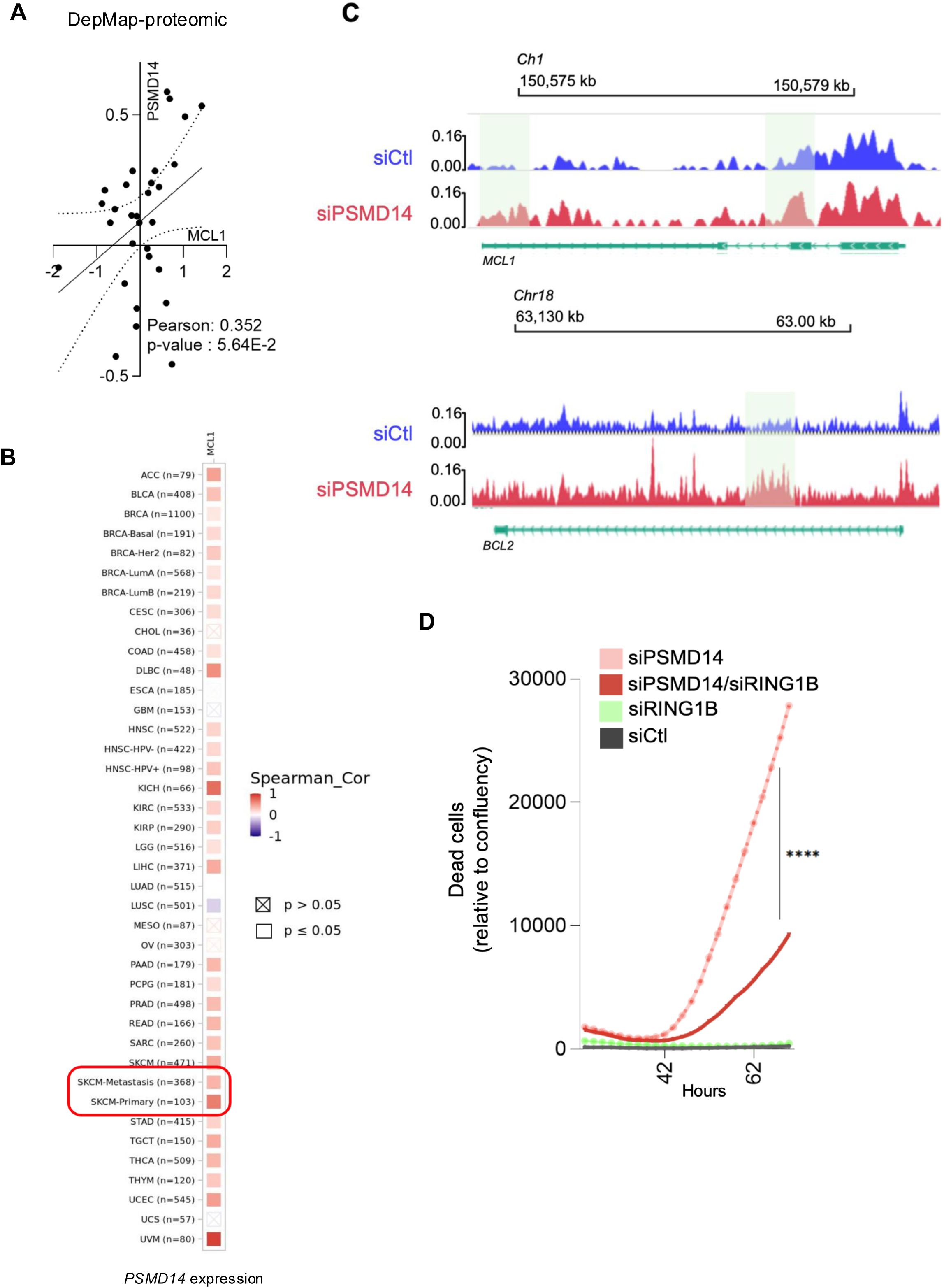
**(A)** Scatter plot with linear regression shows significantly correlated PSMD14 and MCL-1 proteins levels using DepMap-proteomic from a collection of 30 metastatic melanoma cell lines. **(B)** Spearman correlation matrix showing the association between PSMD14 and MCL1 mRNA expression in the TCGA SKCM dataset. **(C)** Representative genomic tracks of H2AK119ub enrichment at the *MCL1* (chr1) and *BCL2* (chr18) loci. ChIP-seq signals for H2AK119ub are shown for cells transfected with a control siRNA (siCtl, blue) or a siRNA targeting PSMD14 (siPSMD14, red). The displayed genomic regions are aligned with the corresponding gene annotations. Shaded green areas highlight regions with notable modulation of H2AK119ub levels. **(D)** Time lapse monitoring of cell death after transfection of A375 cells with control siRNA (siCtl), siPSMD14, siRING1B or the combination of siPSMD14 and siRING1B. Quantification of dead cells is determined using Incucyte® Cytotox red dye relative to cell confluency. Data are the mean ± SEM; ****P < 0.0001, two-way ANOVA test.

**Supplementary Figure 4.**
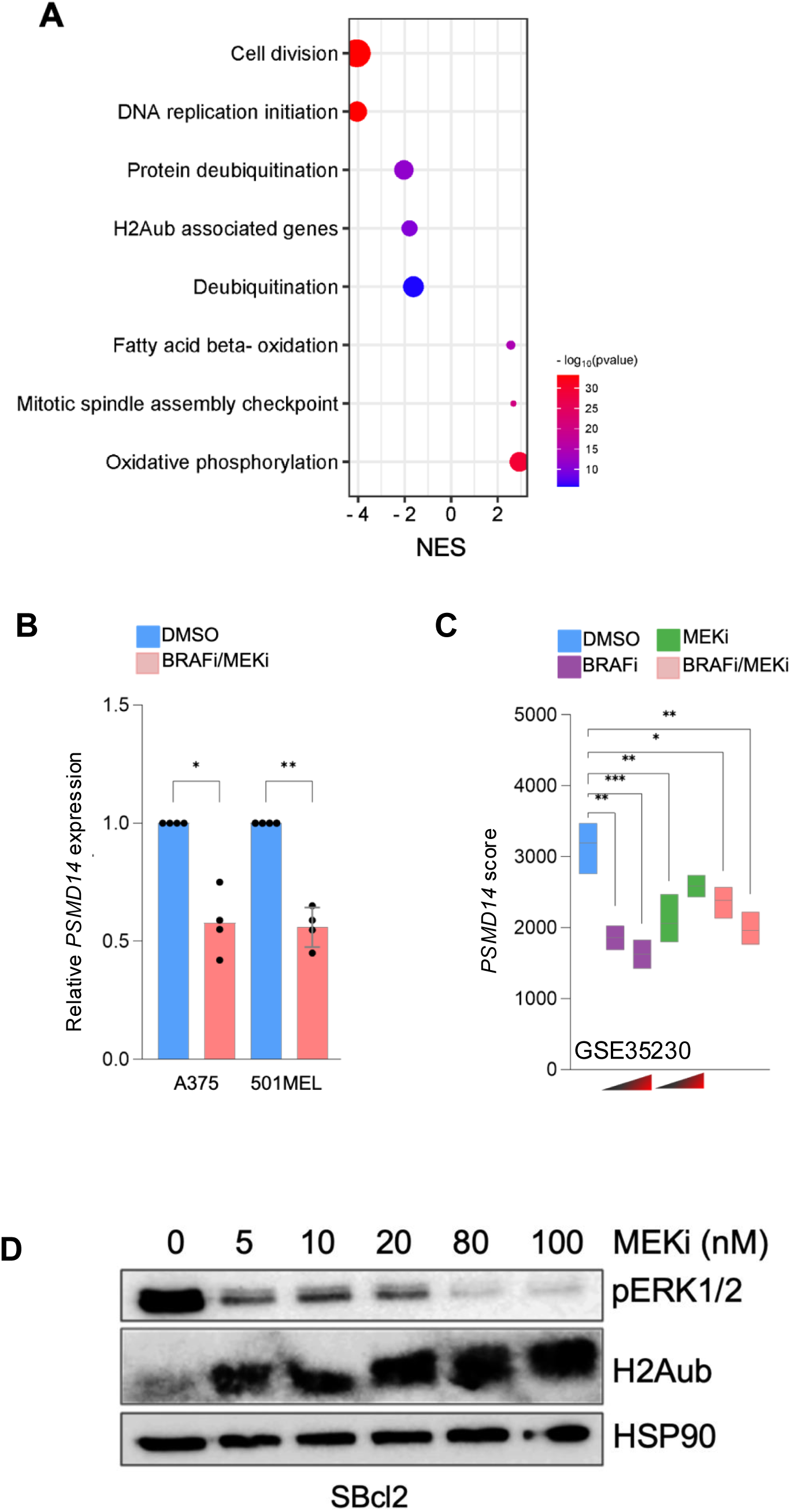
**(A)** Bubble plot showing Gene Ontology (GO) normalized enrichment scores (NES) in BRAFV600E melanoma cells treated with BRAFi versus DMSO (GSE98314). GO terms related to protein deubiquitination and H2Aub target genes are highlighted. FDR-adjusted *p*-values are represented as –log_10_ color scale. **(B)** *PSMD14* expression in A375 cells after BRAF inhibitor (GSK2118436) or MEK inhibitor (GSK1120212) administration alone or in combination (GSE35230). *P ≤ 0.05, **P ≤ 0.01, ***P ≤ 0.001. Two-way ANOVA. **(C)** PSMD14 expression in A375 and 501MEL melanoma cell lines exposed to DMSO or the combination of vemurafenib (10µM) and cobimetinib (1µM) for 24h. Data are presented as mean ± SEM (n= 4). *P < 0.05, **P < 0.01, one way ANOVA. **(D)** Western blot analysis of pERK1/2 and H2Aub levels on NRAS mutant melanoma cells SBcl2 exposed to increasing dose of trametinib (MEKi) for 24h.

**Supplementary Table 1.**
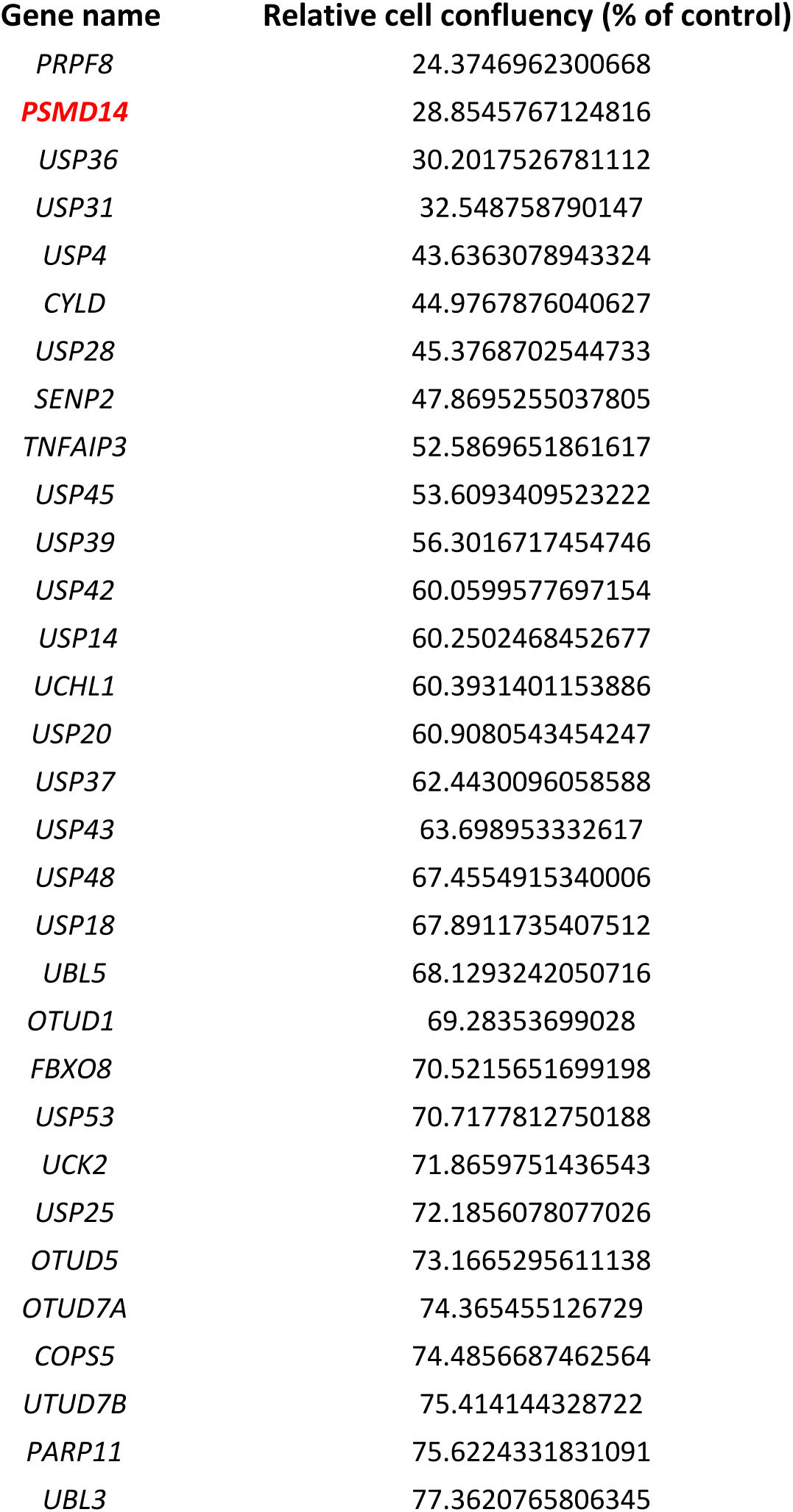
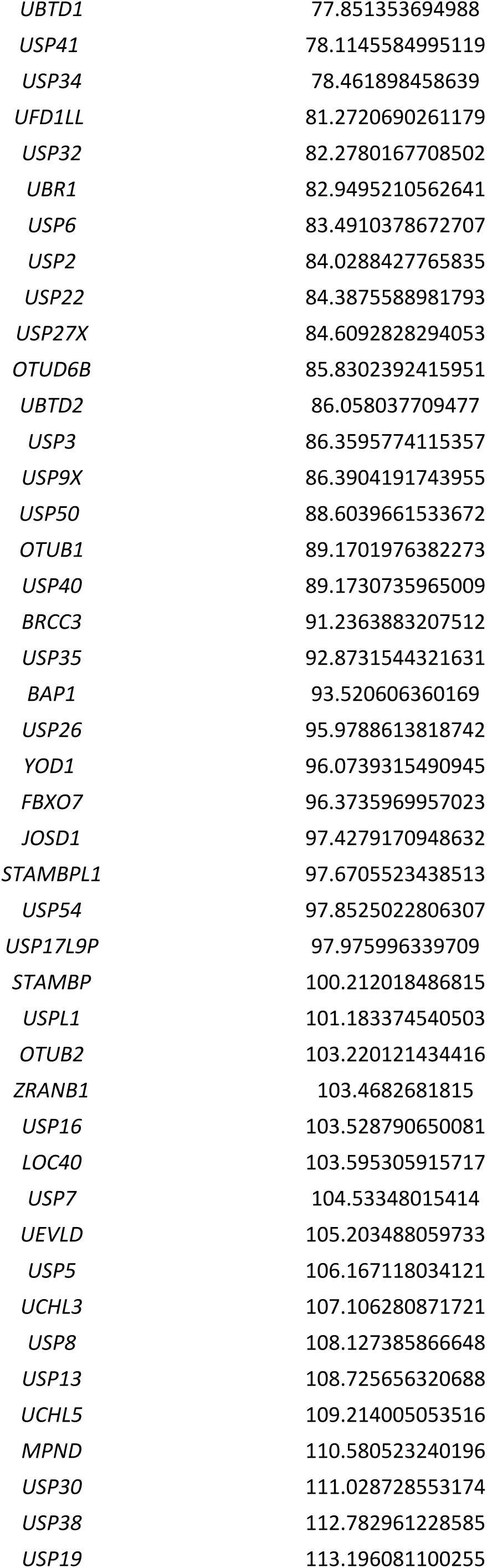
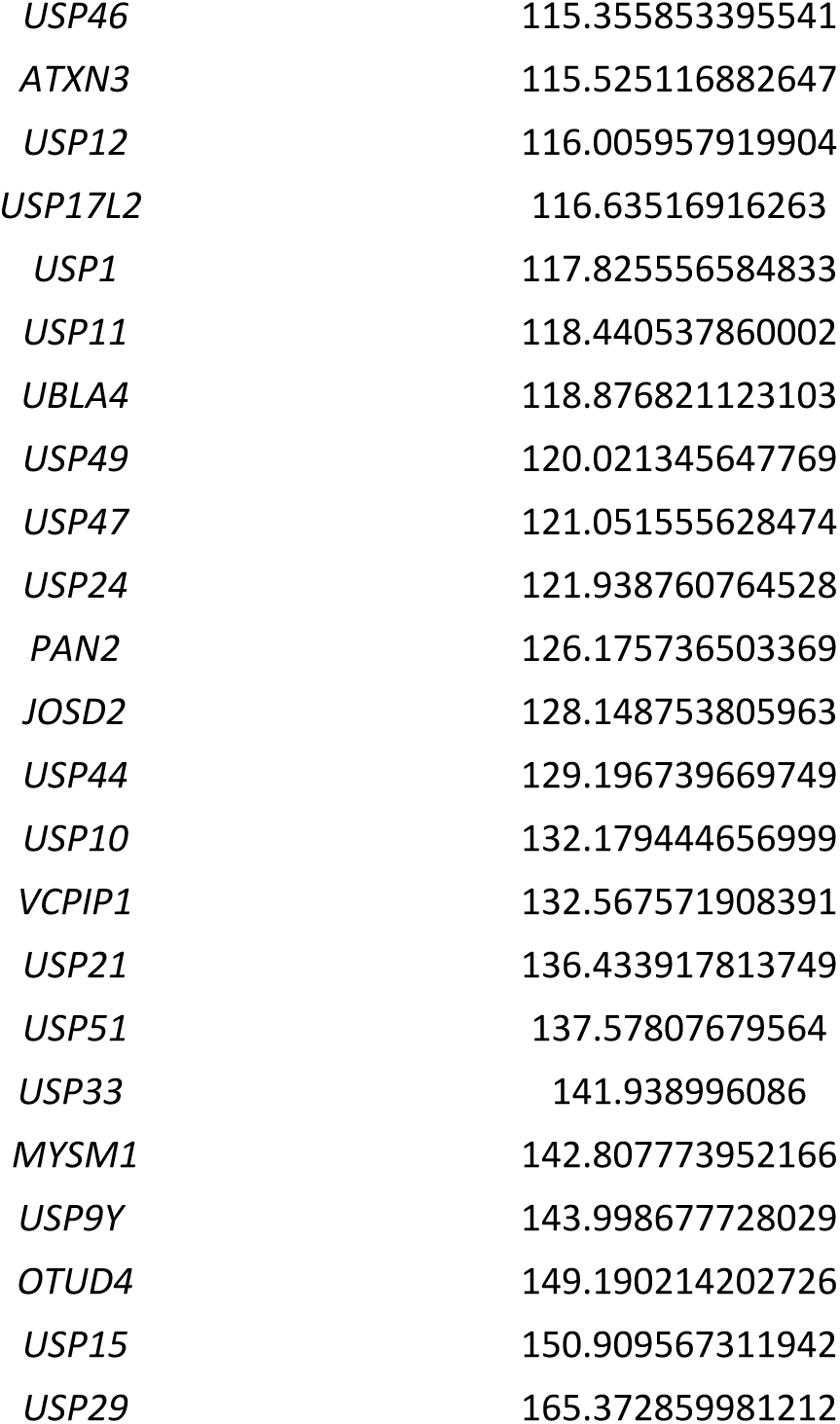
siGENOME DUBs screening: A human DUB siRNA library consisting of pools of four oligonucleotide sequences targeting human DUBs and non-targeting siRNAs were transfected into 501Mel cells and cell proliferation was monitored by live-cell imaging of cell confluency using the IncuCyte ZOOM™ system for 96 h. Cell confluency is represented relative to control non-targeting siRNAs.

**Supplementary Table 2.**
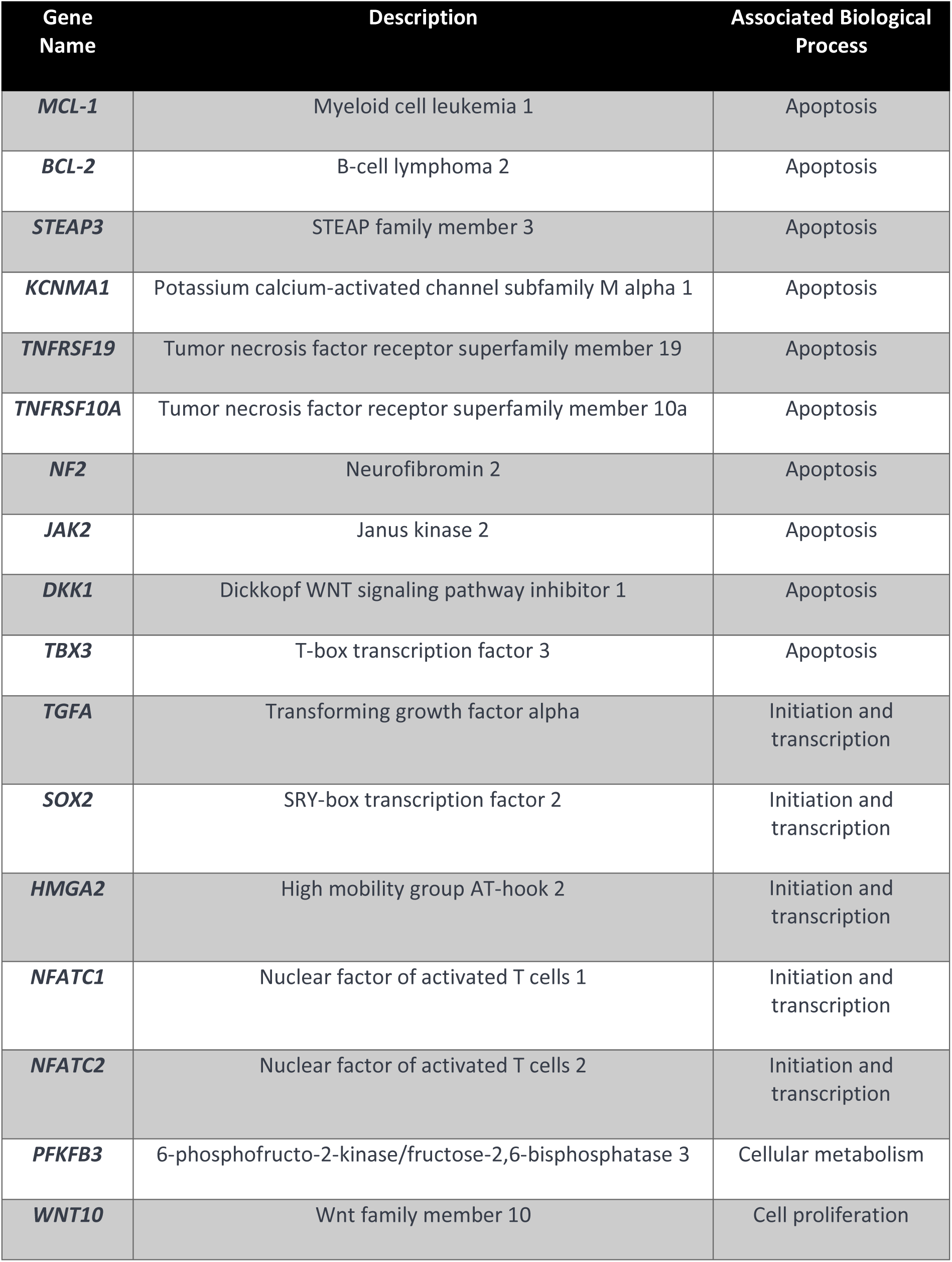
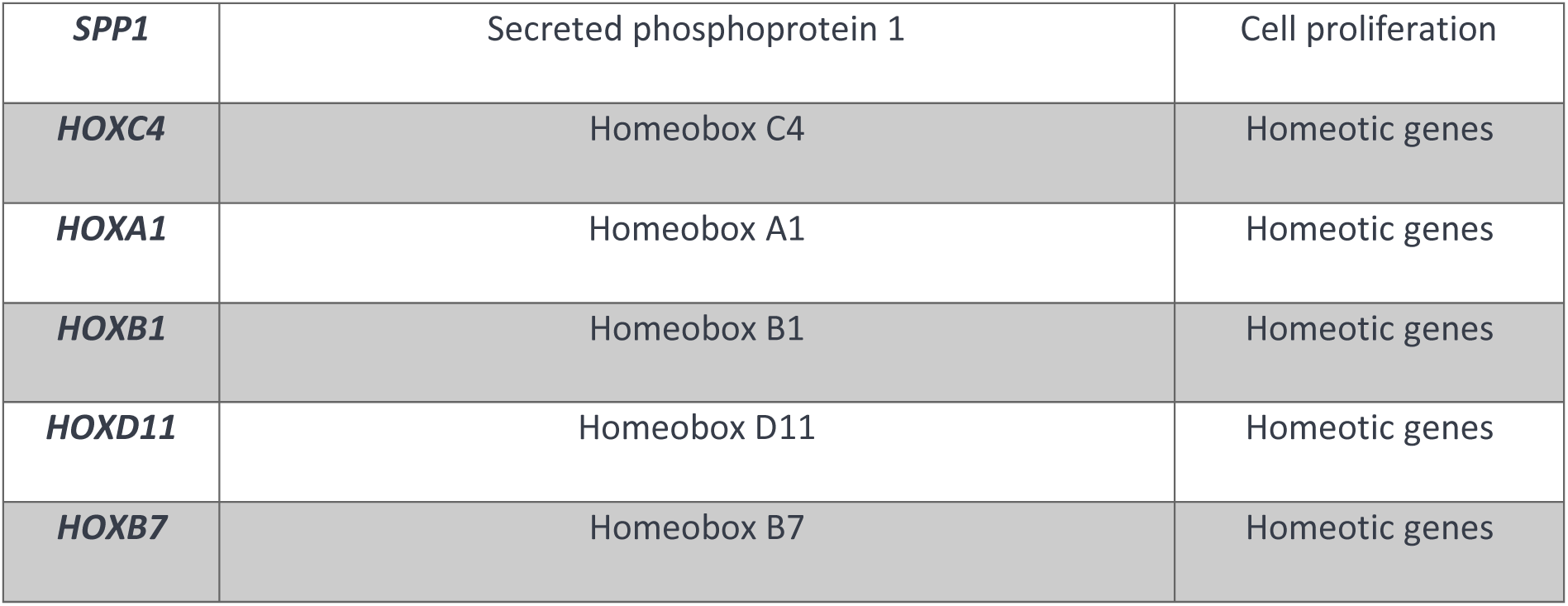
H2Aub gene signature: Selection of genes from the studies by *Zhang et al., 2017* and *He et al., 2019*. These genes were identified enriched in ubiquitinated histone H2A (H2Aub) through ChIP and exhibit decreased expression in response to this modification.

| Gene Name | Description | Associated Biological Process |
| --- | --- | --- |
| ***MCL-1*** | Myeloid cell leukemia 1 | Apoptosis |
| ***BCL-2*** | B-cell lymphoma 2 | Apoptosis |
| ***STEAP3*** | STEAP family member 3 | Apoptosis |
| ***KCNMA1*** | Potassium calcium-activated channel subfamily M alpha 1 | Apoptosis |
| ***TNFRSF19*** | Tumor necrosis factor receptor superfamily member 19 | Apoptosis |
| ***TNFRSF10A*** | Tumor necrosis factor receptor superfamily member 10a | Apoptosis |
| ***NF2*** | Neurofibromin 2 | Apoptosis |
| ***JAK2*** | Janus kinase 2 | Apoptosis |
| ***DKK1*** | Dickkopf WNT signaling pathway inhibitor 1 | Apoptosis |
| ***TBX3*** | T-box transcription factor 3 | Apoptosis |
| ***TGFA*** | Transforming growth factor alpha | Initiation and transcription |
| ***SOX2*** | SRY-box transcription factor 2 | Initiation and transcription |
| ***HMGA2*** | High mobility group AT-hook 2 | Initiation and transcription |
| ***NFATC1*** | Nuclear factor of activated T cells 1 | Initiation and transcription |
| ***NFATC2*** | Nuclear factor of activated T cells 2 | Initiation and transcription |
| ***PFKFB3*** | 6-phosphofructo-2-kinase/fructose-2,6-bisphosphatase 3 | Cellular metabolism |
| ***WNT10*** | Wnt family member 10 | Cell proliferation |
| ***SPP1*** | Secreted phosphoprotein 1 | Cell proliferation |
| ***HOXC4*** | Homeobox C4 | Homeotic genes |
| ***HOXA1*** | Homeobox A1 | Homeotic genes |
| ***HOXB1*** | Homeobox B1 | Homeotic genes |
| ***HOXD11*** | Homeobox D11 | Homeotic genes |
| ***HOXB7*** | Homeobox B7 | Homeotic genes |

## Notes

### Competing Interest Statement

The authors have declared no competing interest.

